# Genomic meta-analysis of the interplay between 3D chromatin organization and gene expression programs under basal and stress conditions

**DOI:** 10.1101/337766

**Authors:** Idan Nurick, Ron Shamir, Ran Elkon

## Abstract

**Background:** Our appreciation of the critical role of the 3D organization of the genome in gene regulation is steadily increasing. Recent 3C-based deep sequencing techniques elucidated a hierarchy of structures that underlie the spatial organization of the genome in the nucleus. At the top of this hierarchical organization are chromosomal territories and the megabase-scale A/B compartments that correlate with transcriptional activity within cells. Below them are the relatively cell-type invariant topologically associated domains (TADs), characterized by high frequency of physical contacts between loci within the same TAD and are assumed to function as regulatory units. Within TADs, chromatin loops bring enhancers and target promoters to close spatial proximity. Yet, we still have only rudimentary understanding how differences in chromatin organization between different cell types affect cell-type specific gene expression programs that are executed under basal and challenged conditions.

**Results:** Here, we carried out a large-scale meta-analysis that integrated Hi-C data from thirteen different cell lines and dozens of ChIP-seq and RNA-seq datasets measured on these cells, either under basal conditions or after treatment. Pairwise comparisons between cell lines demonstrated the strong association between modulation of A/B compartmentalization, differential gene expression and transcription factor (TF) binding events. Furthermore, integrating the analysis of transcriptomes of different cell lines in response to various challenges, we show that 3D organization of cells under basal conditions constrains not only gene expression programs and TF binding profiles that are active under the basal condition but also those induced in response to treatment.

**Conclusions:** Our results further elucidate the role of dynamic genome organization in regulation of differential gene expression between different cell types, and indicate the impact of intra-TAD enhancer-promoter interactions that are established under basal conditions on both the basal and treatment-induced gene expression programs.

## Background

3C-based methods measure the frequency of physical interactions between any pair of genomic loci as a proxy for their spatial proximity. These novel technologies are shedding light on the principles of 3D organization of the genome in the nucleus and its relation to gene regulation [1-3]. A four-layer hierarchy of structures is emerging from these studies [4, 5]. At the top of this hierarchy are the chromosomes which are generally organized in a way that gene-dense chromosomes tend to be at the nuclear interior whereas the more gene-poor chromosomes are found near the nuclear periphery [6]. In the next layer are megabase-scale genomic compartments that are either euchromatic, gene-rich, and transcriptionally active (called A compartments) or heterochromatic, gene-poor, and transcriptionally silent (called B compartments) [5, 7]. Spatially, the open (A-type) compartments cluster together in the nuclear interior, whereas the closed (B-type) compartments tend to cluster near the nuclear periphery [4]. These chromosomal compartments contain ˜100kb-1Mb scale subunits called topologically associating domains (TADs). These are characterized by high frequency of interactions between loci located in the same domain, and much lower interaction rate between loci located in adjacent TADs [8, 9]. Unlike the A/B compartments, which associate to gene expression and therefore markedly vary between different cell types, TADs are largely invariant across different cell types and physiological conditions [7, 10]. At the bottom of the hierarchy are ˜10Kb-1Mb chromatin-looping interactions, bringing enhancers (E) and promoters (P) that are located at high distance along the linear DNA sequence to close spatial proximity. Such E-P loops, a portion of which is cell type specific, mostly occur within TADs and unfrequently cross over TAD boundaries [4, 10]. The 3D organization of the genome has a pronounced cell-to-cell stochastic variability, and the snapshots obtained by 3C-based analyses are typically the result of averaging over a large ensemble of cells.

Our understanding of the roles that the 3D organization of the genome plays in gene regulation has markedly increased in recent years. It emerges that TADs serve as fundamental structural and regulatory building blocks of chromosomes that constrain and largely exclude physical interactions between genes and regulatory elements located in different TADs, while providing sufficiently dynamic local environment that is required for the establishment of intra-TAD E-P links [4, 8]. In line with the view of TADs as structural regulatory units, examination of the dynamic changes in genome 3D organization during differentiation of stem cells into six different linages showed that the regions that changed their A/B compartment mostly corresponded to a single or a series of adjacent TADs [11]. In addition, no significant changes in TAD boundaries were detected in a breast cancer cell-line upon treatment with hormone, suggesting that TADs are also invariant under transient cell challenges [12]. Furthermore, this study found a statistically significant, though limited, number of TADs that behaved as discrete regulatory units where the majority of the genes inside them were either coordinately induced or repressed.

Intra-TAD E-P links are required for the implementation of transcriptional programs that establish and maintain cell identity and responses to environmental cues. How these regulatory interactions are modulated in response to transient perturbation is still not well understood. While some studies have shown that gene induction is accompanied by alterations of chromatin interactions and internal restructuring of TADs [12-14], unexpectedly, it was recently observed that the majority of TNF-α responsive enhancers were already in contact with their target promoters before treatment [15]. Given that the transcriptional responses to various stresses show high level of cell-type specificity, these results suggest that intra-TAD interactions that are already in place in each cell type under basal conditions affect the spectrum of genes that are induced upon triggers in each cell type.

Here, we carried out a large-scale meta-analysis, integrating Hi-C data from 13 different cell lines and dozens of ChIP-seq and RNA-seq datasets recorded in the same cellular systems at basal conditions and in response to various treatments, to further elucidate the intricate interplay between the hierarchical 3D organization of the genome and gene regulation.

## Results

### Differences in gene expression between cell lines correlate with A/B compartmentalization

We first defined the higher order organization of the genome into A/B compartments for 13 human cell lines for which Hi-C data are available (**Supplementary Table 1**). We normalized each Hi-C matrix and performed principal component analysis (PCA) for each intra-chromosomal matrix separately (Methods). By definition, the A compartment is gene rich and is broadly associated with active transcription and epigenomic marks of open chromatin, while the B compartment is gene poor and associates with low transcriptional activity and condensed chromatin. Thus, for each chromosome separately, we used gene density to determine if positive or negative values of the PC that represents the A/B compartmentalization corresponds to A compartment. (Centromeric regions were not included in the A/B partitions since no chromatin interactions are identified by Hi-C in these regions.) **Table 1** summarizes the total genomic size and number of genes assigned to the A and B compartments in each cell line. As an example, **Fig. 1A** shows the partition into A/B compartments we obtained for chromosome 1 in the 13 cell lines. On average, 25% of the genome showed assignment to different compartment in pairwise comparisons between cell lines.

**Table 1.** Genomic size and number of genes assigned to A/B compartments.

| Cell line | A Total size (Mbp) | # Genes in A | B Total size (Mbp) | # Genes in B |
| --- | --- | --- | --- | --- |
| **GM12878** | 1,322 | 15,184 | 1,410 | 3,958 |
| **K562** | 1,376 | 15,401 | 1,356 | 3,741 |
| **HUVEC** | 1,382 | 15,116 | 1,350 | 4,022 |
| **HMEC** | 1,317 | 14,593 | 1,415 | 4,543 |
| **NHEK** | 1,433 | 14,864 | 1,300 | 4,276 |
| **IMR90** | 1,310 | 13,569 | 1,423 | 5,577 |
| **T47D** | 1,372 | 14,114 | 1,350 | 4,979 |
| **MCF7** | 1,384 | 15,056 | 1,450 | 4,758 |
| **MCF10** | 1,386 | 15,090 | 1,451 | 4,772 |
| **LNCAP** | 1,433 | 14,112 | 1,299 | 5,021 |
| **PC3** | 1,395 | 13,341 | 1,313 | 5,692 |
| **KBM7** | 1,301 | 14,506 | 1,431 | 4,631 |
| **PrEC** | 1,327 | 13,994 | 1,387 | 5,041 |

**Figure 1.**
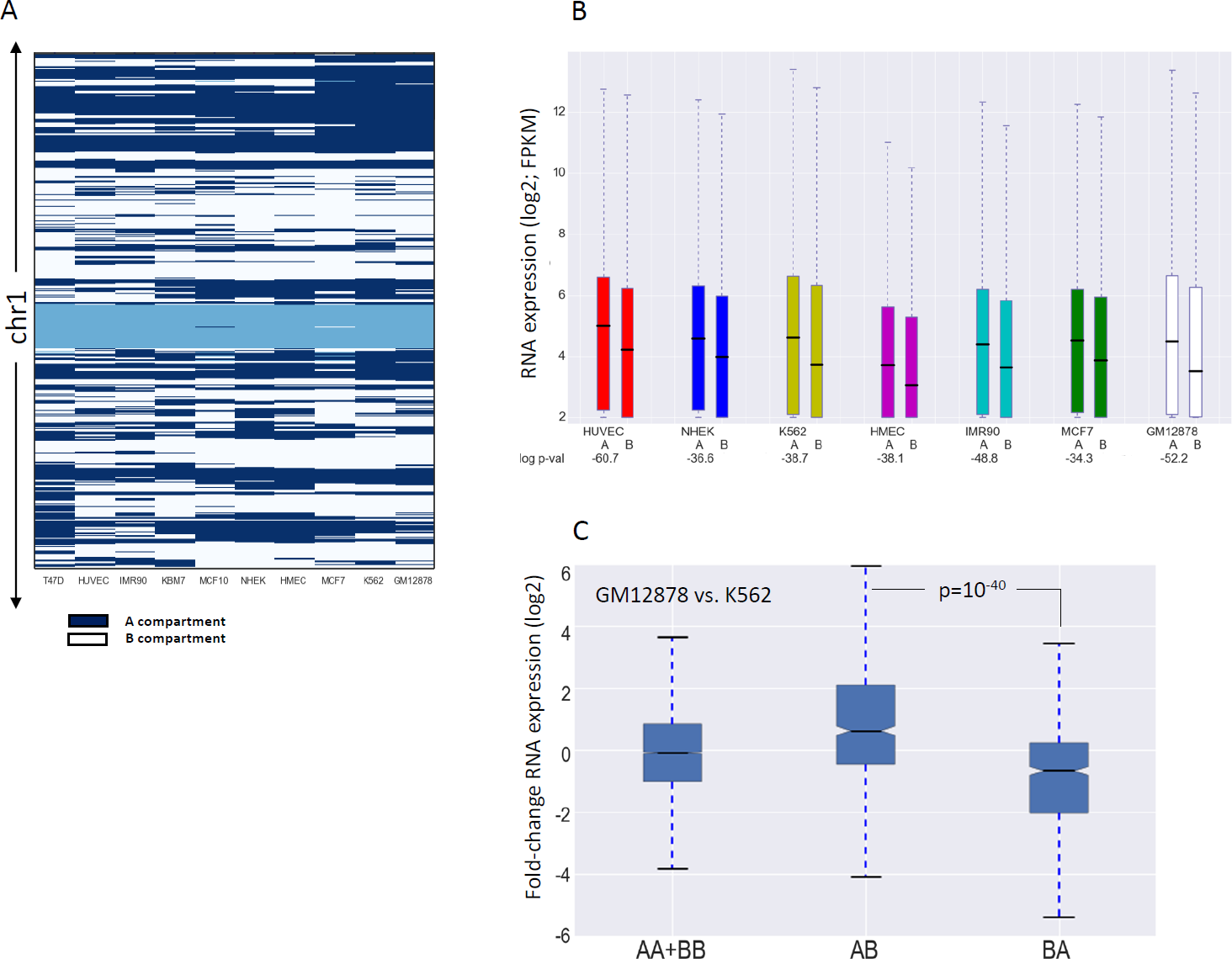
Chromosomal compartments and gene expression. **A**. A/B partition of chromosome 1 for different cell lines based on Hi-C data in 100kb resolution. Dark-blue and white indicate A and B compartments, respectively. Light blue indicates areas which Hi-C could not measure interactions for, e.g., centromeres. **B.** Comparison of gene expression levels in A and B compartments for each cell line. P-values (in log10) for the significance of the difference are indicated below each comparison (Wilcoxon’s test). For all cell lines, genes in A compartment are significantly more highly expressed than genes in B compartment. **C.** Association between dynamic A/B compartmentalization and differential gene expression in the comparison between the GM12878 and K562 cell lines. (AB: the set of genes that are located in compartment A in GM12878 and B in K562; BA: genes located in compartment B in GM12878 and A in K562). Genes in AB have significantly higher expression in GM12878 while genes in BA have higher expression in K562 (p value calculated using Wilcoxon’s test).

As a first examination, per cell line, we confirmed that genes assigned to the A compartment are significantly more highly expressed than genes assigned to the B compartment (**Fig. 1B**). Next, we tested for association between differences in A/B compartmentalization and gene expression across different cell lines. Specifically, for each pair of cell lines, we examined whether genes located in A compartment in one cell line and in B compartment in the other show higher expression in the former. Thus, for each pair of cell lines, we divided the genes into four sets – A in both cell lines (AA), B in both cell lines (BB), A in cell line 1 and B in cell line 2 (AB) and B in cell line 1 and A in cell line 2 (BA). We calculated gene-expression ratios between cell line 1 and 2 and compared the distribution of these ratios between the four gene sets. This analysis confirmed that genes in the AB set are significantly more highly expressed in cell line 1, while genes in the BA set show significantly higher expression in cell line 2 (**Fig. 1C; Fig. S1**).

### Epigenetic differences between cell lines correlate with differences in A/B compartmentalization

As the A compartment is associated with open state of the chromatin we next systematically examined the association between A/B compartmentalization and TF binding. We analyzed 122 TF ChIP-seq datasets recorded by ENCODE for cell lines with Hi-C data (**Supplementary Table 2**). First, we measured TF binding site (TFBS) enrichment for the A compartment, for each cell line separately, by defining the *A-B density factor*, *D* (*D* > 1 implies that binding sites are enriched for the A compartment and *D* < 1 implies that binding sites are enriched for B compartment; Methods). As expected, the chromatin-binding profile of all TFs in all examined cell lines showed a remarkable enrichment for the A compartment (**Fig. S2**; see **Supplementary Table 3** for one detailed example: CTCF).

Next, we examined if A-B transitions between cell-lines are reflected by TF binding profiles. For each pair of cell lines, numbered 1 and 2, we segmented the genome into four regions according to A/B assignment in the two cell lines as described above. For a given TF, we divided the TF binding sites into three groups: sites common to cell line 1 and 2, sites detected only in cell line 1 and sites detected only in cell line 2. We then tested for a relationship between these two divisions. Specifically, we defined the *A-B occupancy enrichment ratio R* (see Methods) to test whether cell-type specific TFBSs occur more often in regions assigned as A compartment in the cell line where the binding occurs and as B in the other cell line than the opposite regions (that is, regions assigned as B-type in the cell line where the binding occurs and as A in the other one). **Table 2A** shows, as an example, the results obtained for CTCF binding sites in the comparison between the HMEC and HUVEC cell lines. As expected, we observed that CTCF BSs specific to HMEC (HUVEC) showed significant preference to AB (BA) genomic regions over BA (AB) regions. As CTCF ChIP-seq data were available for six cell lines with Hi-C data, we could systematically carry out this comparison for this factor. In all pairwise tests, we observed a highly significant preference of CTCF cell-type specific binding to cell-type specific A over B regions (**Fig. 2A**). Yet, a large portion of cell-type specific TFBSs were located in genomic regions that are assigned to A compartment in both cell lines (AA regions) (**Table 2A**), indicating that other factors in addition to A/B compartmentalization determine the TF-chromatin interaction profile in each cell type.

**Table 2.** Comparison of binding site and epigenetic mark occupancy in A/B compartments between two cell types **A.CTCF** **B. H3K9ac** **C. H3K27me3**

| HMEC-HUVEC | AA | AB | BA | BB | total | R | p-value |
| --- | --- | --- | --- | --- | --- | --- | --- |
| HMEC only CTCF BSs | 7,241 | **1,655** | 947 | 4,775 | 14,618 | 2.34 | $10^{-168}$ |
| HUVEC only CTCF BSs | 4,516 | 264 | **1,180** | 1,524 | 7,484 |  |  |
| Common CTCF BSs | 24,986 | 2,587 | 4,647 | 9,750 | 41,970 |  |  |

**B. H3K9ac**
| GM12878 - NHEK | AA | AB | BA | BB | total | R | p-value |
| --- | --- | --- | --- | --- | --- | --- | --- |
| GM12878 only H3K9ac peaks | 14,695 | **3,111** | 596 | 1,708 | 20,110 | 4.54 | $<10^{-300}$ |
| NHEK only H3K9ac peaks | 19,997 | 1,401 | **5,949** | 4,154 | 31,501 |  |  |
| Common H3K9ac peaks | 21,594 | 2,036 | 1,078 | 2,911 | 27,619 |  |  |

**C. H3K27me3**
| MCF7-GM12878 | AA | AB | BA | BB | total | R | p-value |
| --- | --- | --- | --- | --- | --- | --- | --- |
| MCF7 only H3K27me3 peaks | 5,213 | 1,751 | **3,637** | 9,712 | 20,313 | 0.54 | $10^{-122}$ |
| GM12878 only H3K27me3 peaks | 7,176 | **1,765** | 1,145 | 3,608 | 13,694 |  |  |
| Common H3K27me3 peaks | 318 | 95 | 118 | 317 | 848 |  |  |

**Figure 2.**
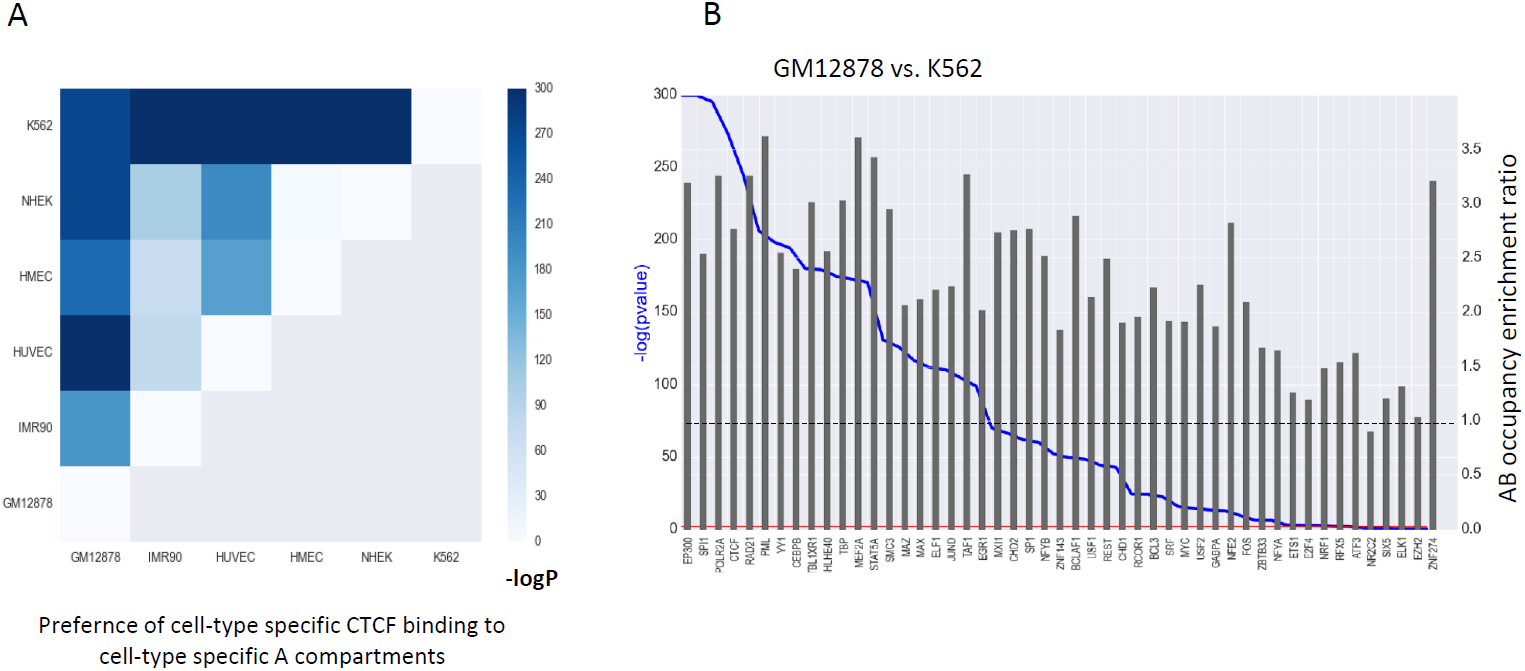
Cell-type specific TF binding vs. cell-type specific compartments. **A**. Relation between cell-type specific A/B partition and CTCF binding sites for six cell lines. For all pair-wise comparisons, CTCF BSs specific to cell 1 (cell 2) showed significant preference to AB (BA) genomic regions over BA (AB) regions. All p-values are highly significant (- log10, chi-square test). **B.** Relationship between cell-type specific TFBSs and A/B compartments in the comparison between GM12878 and K562. For each of the 49 TFs we calculated the *AB occupancy enrichment ratio* as a measure for the preference of its cell-type specific binding events to AB genomic regions over BA regions.

To study the relation between cell-type specific binding sites and compartments across many TFs, we focused on GM12878 and K562, which have ChIP-seq data for 49 common TFs. Strong cell-type specific TFBS-compartment relationship was observed for the vast majority of TFs (**Fig. 2B**). We obtained a significant relationship for 44 out of 49 TFs (FDR < 0.05). (Three out of the five TFs with non-significant p-value have very small group sizes and thus their tests lack statistical power.) The strongest effect was observed for EP300, a transcriptional activator that marks active enhancers.

Next, we carried out similar tests for selected epigenetic marks. H3K9ac, which marks transcriptionally active regions, showed the same correlation between cell-type specific signal and compartmentalization (**Table 2B; Supplementary Table 4A**). Notably, the opposite trend was observed for H3K27me3, which is an epigenetic mark of transcriptionally silenced regions. Namely, in pairwise comparisons between cell lines, regions that showed H3K27me3 signal in only cell line 1 were preferentially associated with BA regions (that is, regions assigned to B compartment in cell line1 and A – in cell line 2) over AB regions (**Table 2C; Supplementary Table 4B**).

### Association between extent of promoter interactions and basal gene expression

The A compartment is generally characterized by high transcriptional activity. Yet, genes within this compartment show considerable expression variability and many of them are not expressed at any detectable level. Our next analysis thus focused on genes within the A compartment, and examined the relationship between the extent of chromatin interactions at promoter regions and gene expression level. In this analysis, we used promoter-enhancer interactions inferred from Hi-C data by the PSYCHIC tool [16]. We expected that, per cell type, promoters of highly expressed genes would show stronger engagement in chromatin interactions than promoters of lowly expressed genes. Indeed, in all five cell lines that we tested, we found a significant positive association between the number of interactions in which a promoter is involved and the gene’s expression level (**Fig. 3A-B; Fig. S3A-B**). We next applied a similar test, but this time using experimental promoter interactions derived from ENCODE’s ChIA-PET data for RNA polymerase II in three cell lines (K562, GM12878 and MCF7). Here too, for all three cell lines examined, we a found a highly significant positive association between extent of promoter interactions and gene expression level (**Fig. 3C-D**; **Fig. S3C-D**).

**Figure 3.**
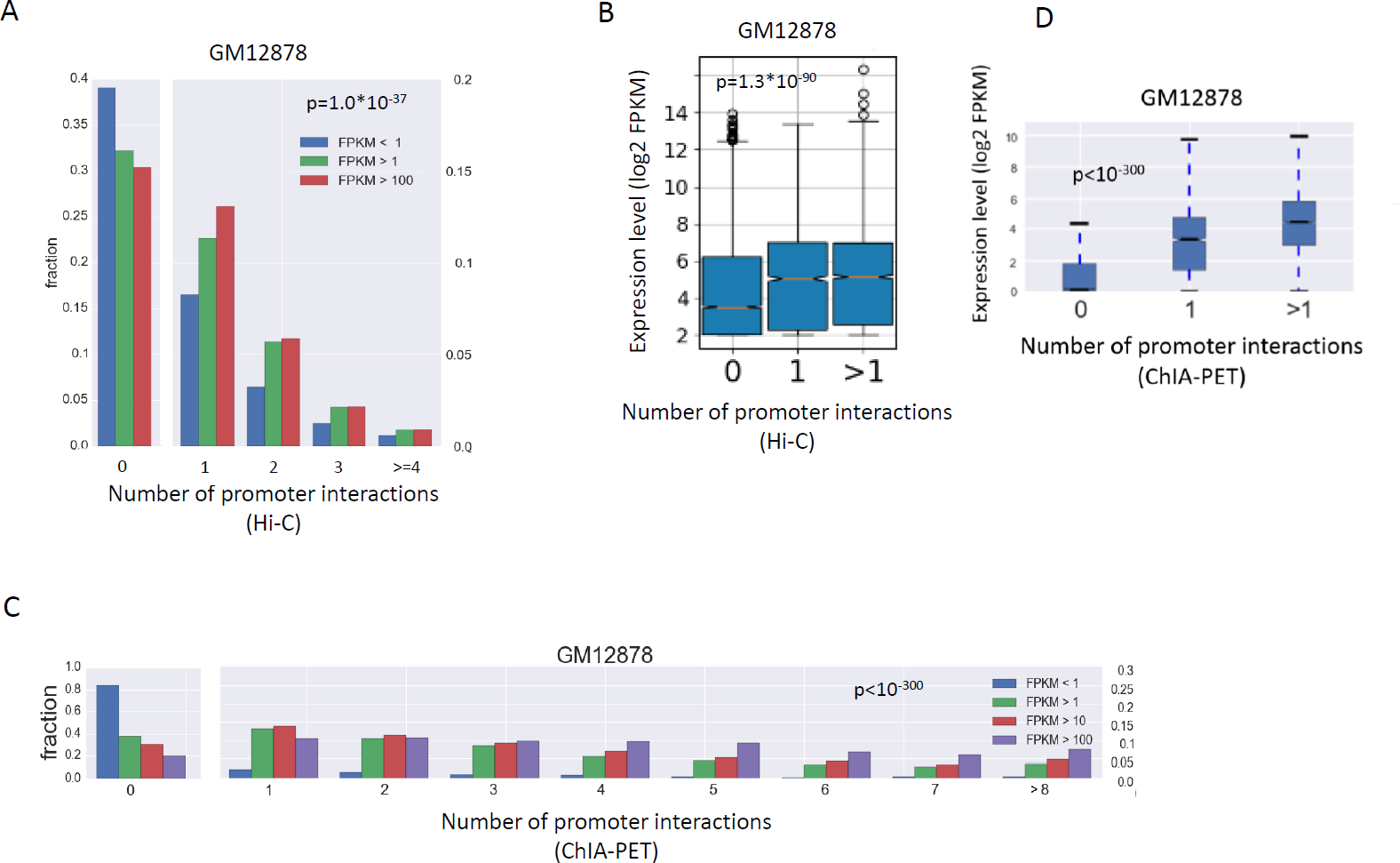
Gene expression levels vs. promoter interactions in compartment A. **A**. Genes in compartment A were partitioned into three groups according to their expression levels. For each group, the distribution of genes over bins of number of promoter interactions, inferred from Hi-C data, is shown. P-value was calculated using Wilcoxon’s test comparing the distributions in the least and most abundant expression groups. **B.** Genes in compartment A were partitioned into three groups according to the number of interactions their promoters are engaged in, and the distributions of gene expression levels were compared (p-value is for Wilcoxon test comparing the group of genes with 0 interactions to the genes with at least one interaction). **C-D.** Same analysis as in B and A, respectively, but here using promoter interactions derived from RNA PolII ChIA-PET data in the GM12878 cell line.

The above analysis was done on each cell line separately. We next examined correlation between dynamic promoter interactions and gene expression across cell lines. Specifically, we tested if changes in a gene’s expression over different cell lines are associated with differences in the number of interactions involving the gene’s promoter in these cell lines. This analysis too was confined to genes located within the A compartment in both cell lines (“AA” genes). For each pair of cell lines, we divided the genes into four groups, based on RNA Pol-II ChIA-PET data: no promoter interactions in both cells (“00” group); promoter interactions detected only in cell line 1 (“10” genes); only in cell line 2 (“01” genes) and in both (“11” genes). Notably, differential gene expression between pairs of cell lines was strongly associated with differential engagement of promoters in chromatin interactions (**Fig. 4A-B** for MCF7 vs. K562). Similar results were obtained for the other pairs that we examined (data not shown). These results indicate that dynamic, intra TAD chromatin interactions involving gene promoters within the A compartment modulate cell-type specific gene expression.

**Figure 4.**
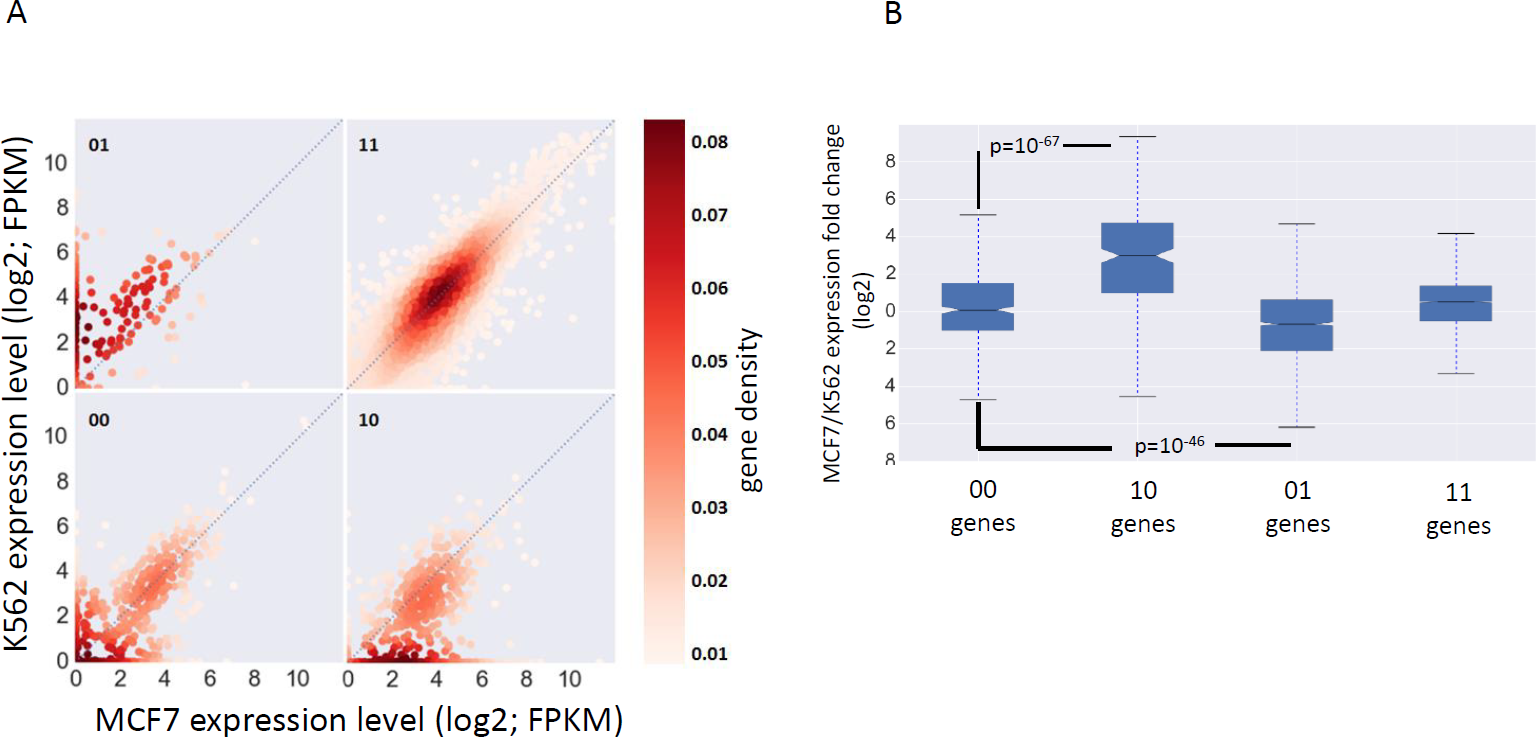
Differential promoter interactions vs. differential gene expression. **A**. Genes located in the A compartment in both MCF7 and K562 cell lines (AA genes) were divided into four groups according to the engagement of their promoters in chromatin interactions in the two cell lines (as indicated by RNA PolII ChIA-PET data). Gene sets 00, 01, 10, and 11 correspond, respectively, to the set of genes whose promoter is engaged in chromatin interactions in either none, only K562, only MCF7 or both cell lines. Expression levels in both cell lines are plotted for each gene set. Color indicates gene density. **B**. Distribution of fold-change in gene expression (log2) between MCF7 and K562 was calculated for each gene set. Highly significant association between differential gene expression and differential involvement of promoters in chromatin interaction was observed. p-values are computed using Wilcoxon test.

### Association between basal chromatin organization and treatment-induced TF binding profiles

Many transcriptomic studies demonstrated that a large portion of the transcriptional response to various challenges is cell-type specific [17-19]. Surprisingly, recent epigenomic and transcriptomic analysis of the response to TNFα observed that enhancers activated by this trigger were already in contact with their target promoters before treatment [15]. Therefore, we next sought to examine the role of basal chromatin interactions, which are in place in cells before any challenge is applied, in shaping cell-type responses induced by treatment. To allow us to draw general conclusions, we analyzed a variety of cell lines and multiple treatments covering diverse biological processes. We first analyzed 110 publicly available ChIP-seq datasets, recorded in cells for which we analyzed Hi-C data, that profiled TF binding and epigenetic marks before and after the application of various treatments. Overall, we analyzed 21 TFs in seven cell lines in response to 22 treatments. Per experiment, we analyzed TFBSs detected under basal and stress conditions and identified the set of TFBSs that were induced in response to treatment. We then divided these induced TFBSs into A/B compartments. For the vast majority of experiments (>90%), we observed a highly significant preference of the induced sites to the A compartment, indicating that the preexisting A/B compartmentalization within a cell line constrains TF-chromatin interactions that are induced in response to stress **(Fig. 5A** and **Supplementary Table** 5).

**Figure 5.**
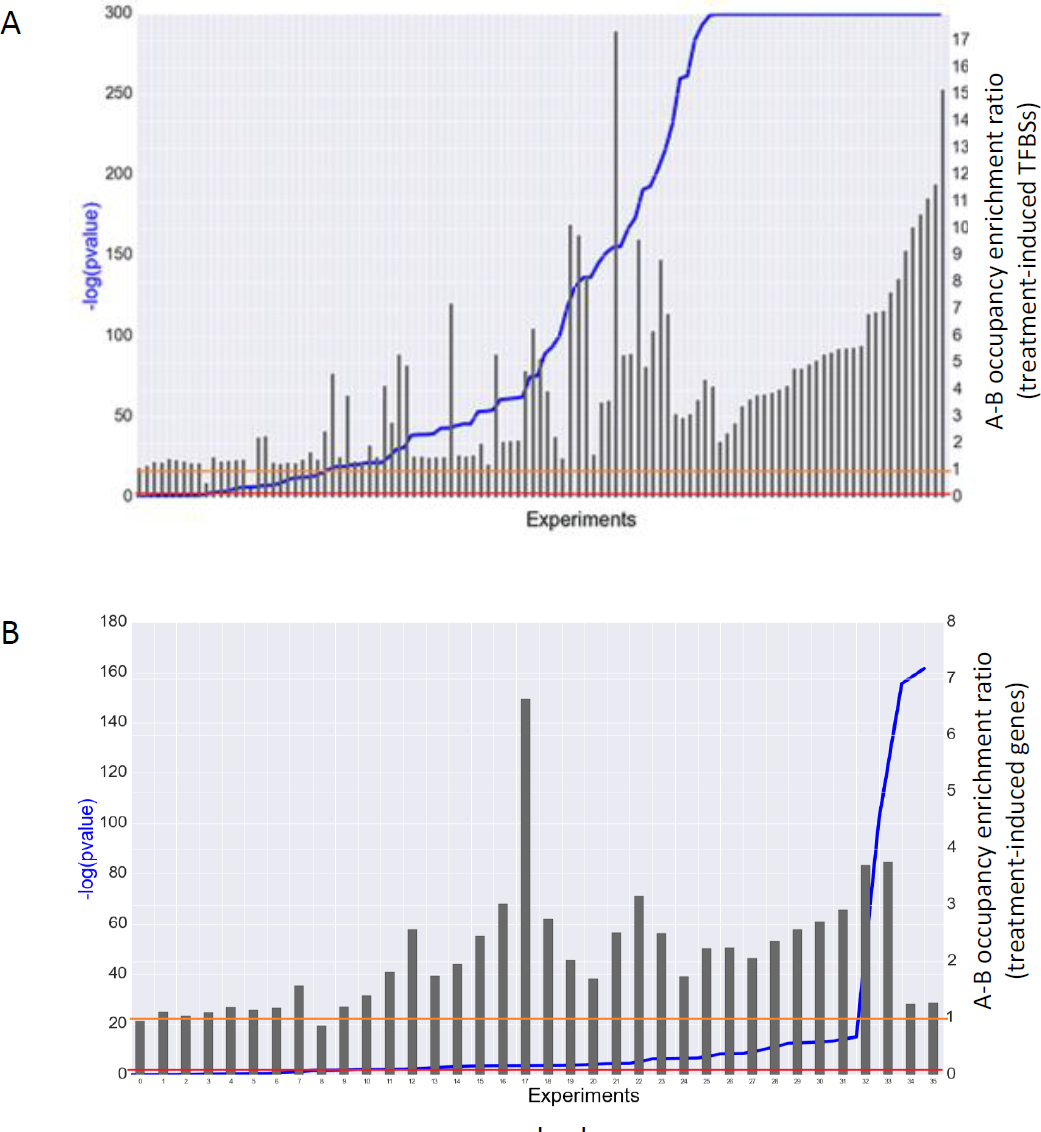
Enrichment of treatment-induced TFBSs and genes in the basal A compartment. **A**. TFBSs induced by various treatments are significantly enriched in the A compartment (as determined in the cells under basal condition). Experiments are sorted by p-value, enrichment ratios are represented by bars. Red line: p-value = 0.01, orange line: enrichment ratio = 1. **B.** Same as A, but for treatment-induced genes.

Next, to examine the relationship between cell-type specific chromatin organization and response to treatment more directly, we sought ChIP-seq datasets that profiled the same TF in response to the same treatment in different cell lines (for which we also analyzed Hi-C data). Several experiments that examined responses to TNFα and estradiol met this requirement. For each pair of cell lines treated by the same agent and profiled for the same TF, we again divided the induced TFBSs into three groups: binding sites induced upon treatment only in cell line 1, binding sites induced only in cell line 2 and binding sites induced in both. Induced TFBSs in each group were then divided into four categories – AA, AB, BA and BB as defined above. In all comparisons, TFBSs induced only in cell line 1 showed significant preference for AB regions over the BA ones, and vice versa for TFBSs induced only in cell line 2 (**Table 3**; **Supplementary Table 6**). This result further demonstrates the impact of cell-specific basal genome organization on the landscape of TF-chromatin interactions that are induced upon challenge. Note that despite the significant association between cell-type specific TF binding induction and chromatin organization, most of the cell-type specific induced TFBSs were located in AA regions (**Table 3**; **Supplementary Table 6**), indicating that factors other than A/B compartmentalization play more dominant role in determining cell-type specific TF binding profiles.

**Table 3.** Cell-type specific treatment-induced TFBSs show preference to cell-type specific A compartment.

| MCF7 - T47D;<br>Treatment: Estradiol<br>Antibody: ER | AA | AB | BA | BB | total | A/B<br>Enrichment | R | p-value |
| --- | --- | --- | --- | --- | --- | --- | --- | --- |
| MCF7 only ESR1 BSs | 9,656 | **2,356** | 1,459 | 3,285 | 16,756 | 1.61 | 1.61 | 3.29E-29 |
| T47D only ESR1 BSs | 2,031 | 257 | **410** | 475 | 3,173 | 1.6 |  |  |
| Common ESR1 BSs | 3,138 | 406 | 330 | 461 | 4,335 |  |  |  |

### Association between basal chromatin organization and transcriptional response to treatment

The above analyses examined the association between basal chromatin 3D organization and treatment-induced TF-chromatin interactions. Next, we examined the association between basal chromatin 3D organization and gene induction in response to treatment. For this goal, we analyzed 36 gene expression datasets that recorded transcriptome profiles in response to various challenges (in cell lines for which we have analyzed Hi-C data). For each cell line and treatment, we tested if the set of genes that were induced upon treatment was over-represented in the A compartment. Indeed, in most conditions, we observed a significant preference of the induced genes to the A compartment (**Fig. 5B** and **Supplementary Table 7**). This suggests that the preference of induced TFBSs to the pre-challenge A compartment leads to an induced transcriptional response that show similar preference. The statistical significance obtained by the analysis of the induced genes was usually lower than that obtained by the induced TFBSs since the numbers of responsive genes were substantially lower than the numbers of induced TFBSs. Nevertheless, 28 out of 34 experiments had a significant p-value (FDR < 0.05) and 32 out of 34 experiments had enrichment factor larger than 1.0 (p < 3.5 ∗ 10^−8^; binomial test).

In a previous section, we described an association between the extent of promoter interactions and basal gene expression level (**Fig. 3**). Here, we examined if promoters of genes, within compartment A, that were induced in response to challenges also show higher involvement in chromatin interactions that already exist in the cells under basal condition. Analyzing numerous RNA-seq datasets, we systematically observed that promoters of induced genes were engaged, already in basal conditions, in a markedly higher number of chromatin interactions compared to promoters of non-induced genes that are located in the A compartment and have comparable basal expression levels. We estimated the significance of this higher degree of chromatin interaction by using a permutation test with 10,000 iterations, in each iteration selecting a random set of genes (from A compartment) of the same size as the induced genes set. Expression level was controlled for by dividing the A-compartment genes into 10 bins, according to their basal expression level, and generating random gene sets having the same distribution as the test set of the induced gene. In all experiments except one (with very low number of included genes and thus limited statistical power), we obtained significant p-values (p<0.05) (**Fig. 6**; **Supplementary Table 8**).

**Figure 6.**
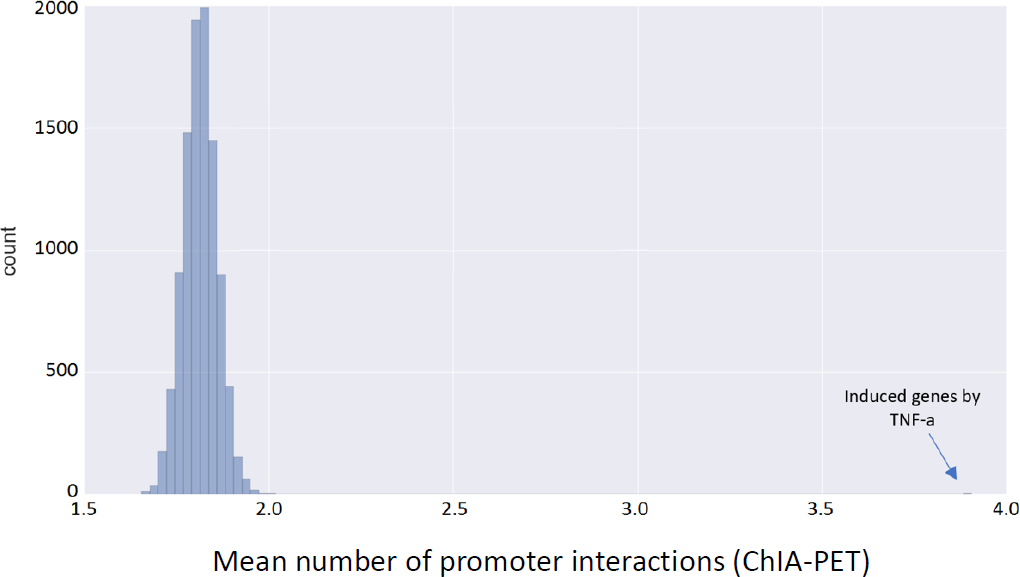
Engagement of promoters of treatment-induced genes in basal chromatin interactions. We used permutation tests to assess the significance of the engagement in chromatin interactions observed for promoters of genes that were induced upon challenges. The figure shows the analysis for the set of genes that were induced in GM12878 cells upon TNFα treatment (the positive set). 10,000 randomly selected gene sets of the same size and with the same basal expression distribution as the positive set were used to generate a null distribution. The mean number of promoter interactions per gene (3.9) was significantly higher for the positive set.

Last, we examined if cell-type specific gene induction in response to treatment correlates with pre-existing chromatin compartmentalization. We focused on response to TNFα as we gathered RNA-seq datasets that profiled responses to this trigger in five different cell lines for which we determined AB compartmentalization based on Hi-C data (HMEC, IMR90, GM12878, MCF7 and HUVEC). We followed the same analysis that we applied above to TFBSs that were induced in a cell-type specific manner (Table 3), and applied it to the set of TNFα-induced genes. For 8 out of 10 pairwise comparisons we found a strong association: genes induced specifically in cell line 1 were significantly enriched for AB over BA regions (and vice versa for genes specifically induced in cell line 2) (**Table 4; Supplementary Table 9**). Notably, in this analysis too, the majority of cell-type specific responsive genes were located in AA regions, again indicating that other factors play critical roles in determining the specific spectrum of genes that respond to a challenge in each cell type.

**Table 4.** Cell-type specific treatment-induced genes show preference to cell-type specific basal A compartment.

| **HMEC - MCF7** | **AA** | **AB** | **BA** | **BB** | **Total** | **R** | **Enrichment** | **p-value** |
| --- | --- | --- | --- | --- | --- | --- | --- | --- |
| **Induced only in HMEC** | 29 | **20** | 6 | 9 | 64 | 3.44 | 2.0 | 0.00095 |
| **Induced only in MCF7** | 267 | 22 | **36** | 33 | 358 | 1.67 |  |  |
| **induced in both** | 34 | 4 | 1 | 4 | 43 |  |  |  |

### Discussion

To further explore links between the 3D organization of the genome and gene regulation, we have analyzed together Hi-C data from 13 different human cell lines and numerous ChIP-seq and RNA-seq experiments that were recorded in the same cell lines under basal conditions and in response to various treatments. We first confirmed the strong relationship between the partition of the genome into the A/B compartments and transcriptional activity. In all cell lines, expression level of genes located within the A compartment was significantly higher than expression level of genes located in B (**Fig. 1B**), and differential expression between cell lines correlated with differences in AB compartmentalization (**Fig. 1C**). Similarly, in analysis of 122 TF ChIP-seq datasets, the binding profile of the vast majority of TFs showed a significant preference for the A compartment (**Fig. S2**), and cell-type specific TF binding events correlated with cell-type specific A compartments (**Fig. 2**). As expected, the H3K27me3 epigenetic modification that marks transcriptionally silenced regions showed the opposite preference and was highly enriched in the B compartment (**Table 2C**). These results demonstrate the effect of higher order chromosomal organization on TF-chromatin interaction profiles. Yet, in comparisons of TF binding profiles between different cell lines, the majority of cell-type specific TFBSs were located in genomic regions that are assigned to A compartment in both cell lines (AA regions) (**Table 2**). This observation indicates that other factors play stronger roles than A/B compartmentalization in shaping the landscape of TF-chromatin interactions in each cell type. Master TFs that establish and maintain cell identify are likely a major factor. These regulators exhibit highly cell-type specific expression pattern and were shown to have great impact on the selection of TF binding sites in different cell types [20, 21].

Using E-P links derived from Hi-C and ChIA-PET data, we found a strong correlation between gene expression levels and the extent to which promoters are engaged in chromatin interactions (**Fig. 3**). Moreover, we showed that differential expression between cell types is associated with dynamic change in involvement of promoters in such interactions (**Fig. 4**), which are most likely mediated by cell type specific TFs. A recent study showed that during cell reprogramming, the expression of lineage-specific TFs drives genome reorganization that precedes changes in gene expression [22].

We then turned to analyze the impact of the organization of the genome under basal conditions on transcriptional programs that are induced in response to various triggers. First, we showed that both induced TF binding events and induced genes are enriched in the A compartment (**Fig. 5**), indicating that preexisting A/B compartments within a cell constrains its network of induced TF-chromatin interactions and activated genes. We then demonstrated the association between cell-type specific response to triggers and basal cell-type specific AB compartmentalization. Cell-type specific induced TF binding and activated genes show significant enrichment for cell-type specific A compartments (**Table 3 and 4**). Yet, here too, a large portion of the cell-type specific induced TFBS and genes are located in genomic regions that are assigned to the A compartment in both responsive and non-responsive cell lines, further underscoring that additional key factors participate in shaping the specific transcriptional response to challenges elicited in each cell type.

Current techniques for determining the 3D organization of the genome are still limited in their resolution and sensitivity. Further development of these methods together with advances in their application to single cells will allow us to better understand how the genome is reorganized at multiple structural layers in response to various triggers and stresses, and to elucidate how this topological reorganization is causally linked to cell-type specific transcriptional programs that are induced in response to these challenges.

### Conclusions

Collectively, the large-scale meta-analysis that we carried out in this study further demonstrates the strong association between cell-type specific A/B compartmentalization, modulation of landscape of TF-chromatin interactions and differential gene expression. Moreover, our results further indicate a role for the 3D organization of the genome under basal conditions, at the layers of both A/B compartmentalization and intra-TAD enhancer-promoter interactions, in shaping TF binding events and the network of genes that are induced in response to treatment.

## Methods

### Identification of A and B compartments from Hi-C data

We defined A/B compartments for 13 human cell lines for which Hi-C data are available (**Supplementary Table 1**). Identification of A and B compartments was performed similarly to what has been previously described [5, 11]. Briefly, Hi-C contact frequency matrix was first normalized using the Knight and Ruiz matrix balancing method [23]. Then, we performed principal component analysis (PCA) for each intrachromosomal matrix separately. In most cases, the first principal component vector partitions the chromosome into two compartments, A and B, according to the sign of the elements. In the other cases, mostly in short chromosomes, the first principal component divides the chromosome to its two arms and the second component partition it to the A/B compartments. As seen in previous studies [7], the A compartment is gene rich and its chromatin is less dense, while the B regions are gene poor and their chromatin is denser. Thus, we determined, for each chromosome separately, whether positive or negative values of the PC that indicates the A/B compartmentalization correspond to A or B based on gene richness; the compartment with higher gene density was labeled as A compartment. Centromeric regions were not included in the A/B partitions since no chromatin interactions are identified by Hi-C in these regions.

### RNA-seq analysis

RNA-seq data were analyzed using a standard pipeline. Briefly, raw sequence data were downloaded from GEO/SRA DB, and mapped to the human genome (hg19) using tophat2 [24]. The number of reads that mapped to each annotated gene was counted using HTSeq-counts [25] based on GENCODE annotations [26]. Gene expression estimates were normalized to RPKM. We compared expression profiles between treated and control samples and defined the genes whose expression was changed by at least 1.5-fold as differential genes. (To avoid inflation of lowly expressed genes among the called differential genes we used a floor level of 1.0 RPKM).

### ChIP-seq analysis

To ensure analysis uniformity, we did not rely on peaks called by original studies, but downloaded raw sequence data and detected TF peaks ourselves. Briefly, for each ChIP-seq experiment, reads were aligned to the human genome (hg19) using bowtie2 [27] and peaks were called using MACS2 by comparing IP and input samples. For detection of peaks induced upon treatment, IP samples measured under control and treated conditions were directly compared [28].

### AB Density factor D

For each transcription factor and cell line we computed the *AB density factor*, *D*, defined as follows: Let the number of observed binding sites in region *S* be *O*(*S*) and number of expected binding sites in region *S* be *E*(*S*):

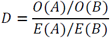

*D* > 1 implies that binding sites are enriched for A compartment and *D* < 1 implies that binding sites are enriched for B compartment. For TF binding sites, *E*(*A*)/*E*(*B*) is equal to the ratio between the genomic size of the two compartments.

### AB Occupancy enrichment ratio R

For pairwise comparisons between cells, to test if cell-type specific TFBSs occur more often in regions assigned as A compartment in the cell line where the binding event was detected and as B in the other cell line than the opposite assignment, we defined the *AB occupancy enrichment ratio* R, as follows: Let the number of BSs in region S occurring only in cell line *i* be *n*(*i*, *S*). Then

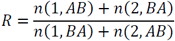

## Declarations

### Ethics approval and consent to participate

Not applicable

### Consent for publication

Not applicable

### Availability of data and material

All analyses carried out in this study used publicly available datasets. References are provided in the supplementary tables.

### Authors’ contributions

I.N., R.S., and R.E. designed the research. I.N. performed the analyses, R.E. and R.S. critically reviewed the analyses and. wrote the manuscript. All authors approved the final version.

## Acknowledgements.

R.E. is a Faculty Fellow of the Edmond J. Safra Center for Bioinformatics at Tel Aviv University.

## Supplementary Figure legends

**Supplementary Table 1.**
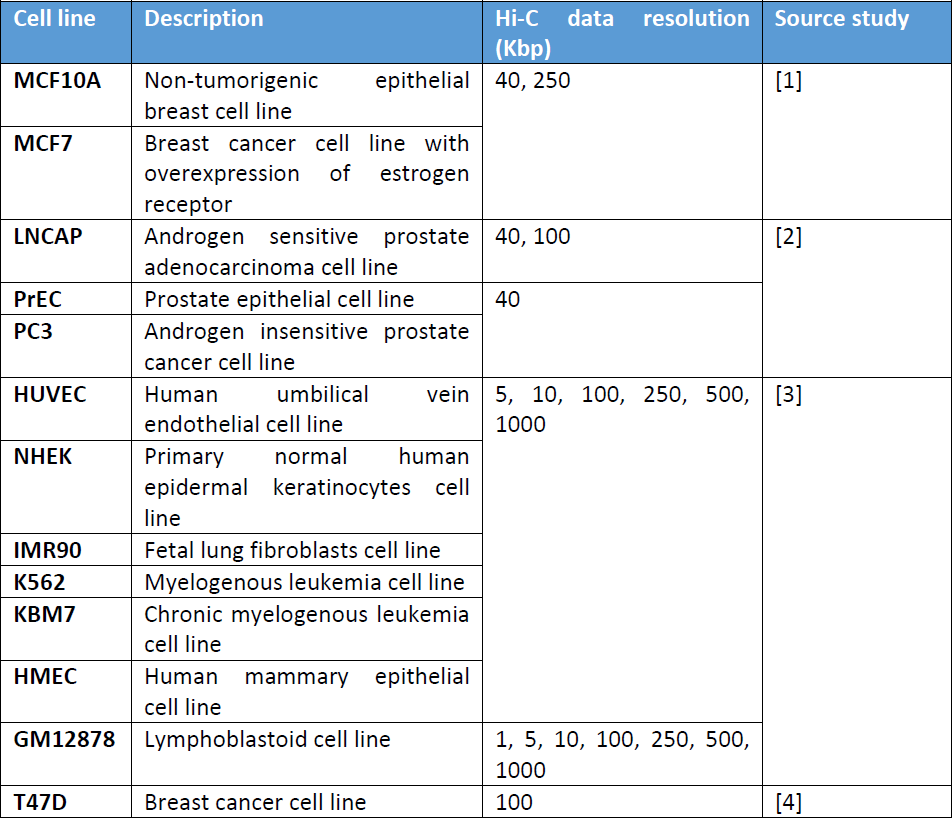
Hi-C datasets

**Supplementary Table 2.**
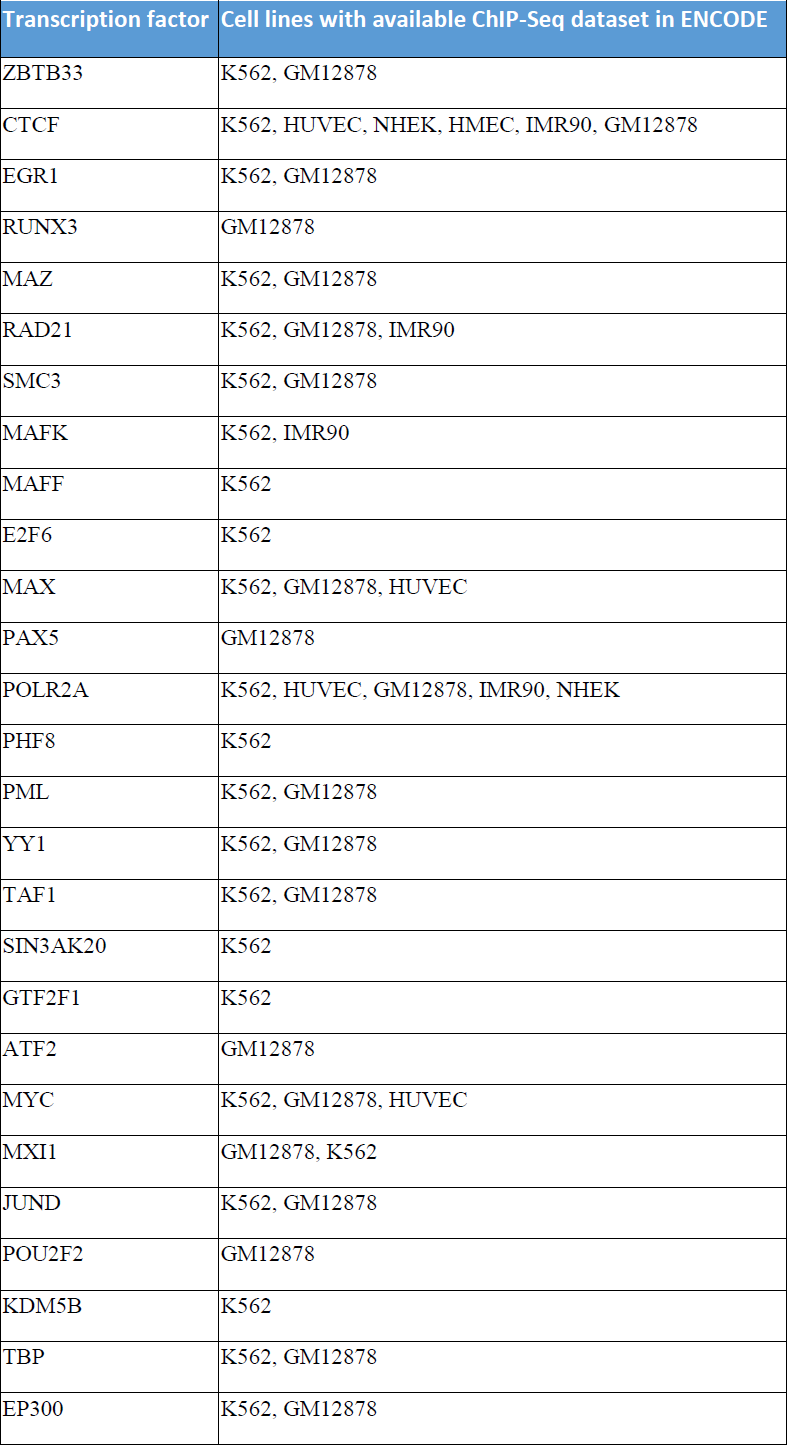
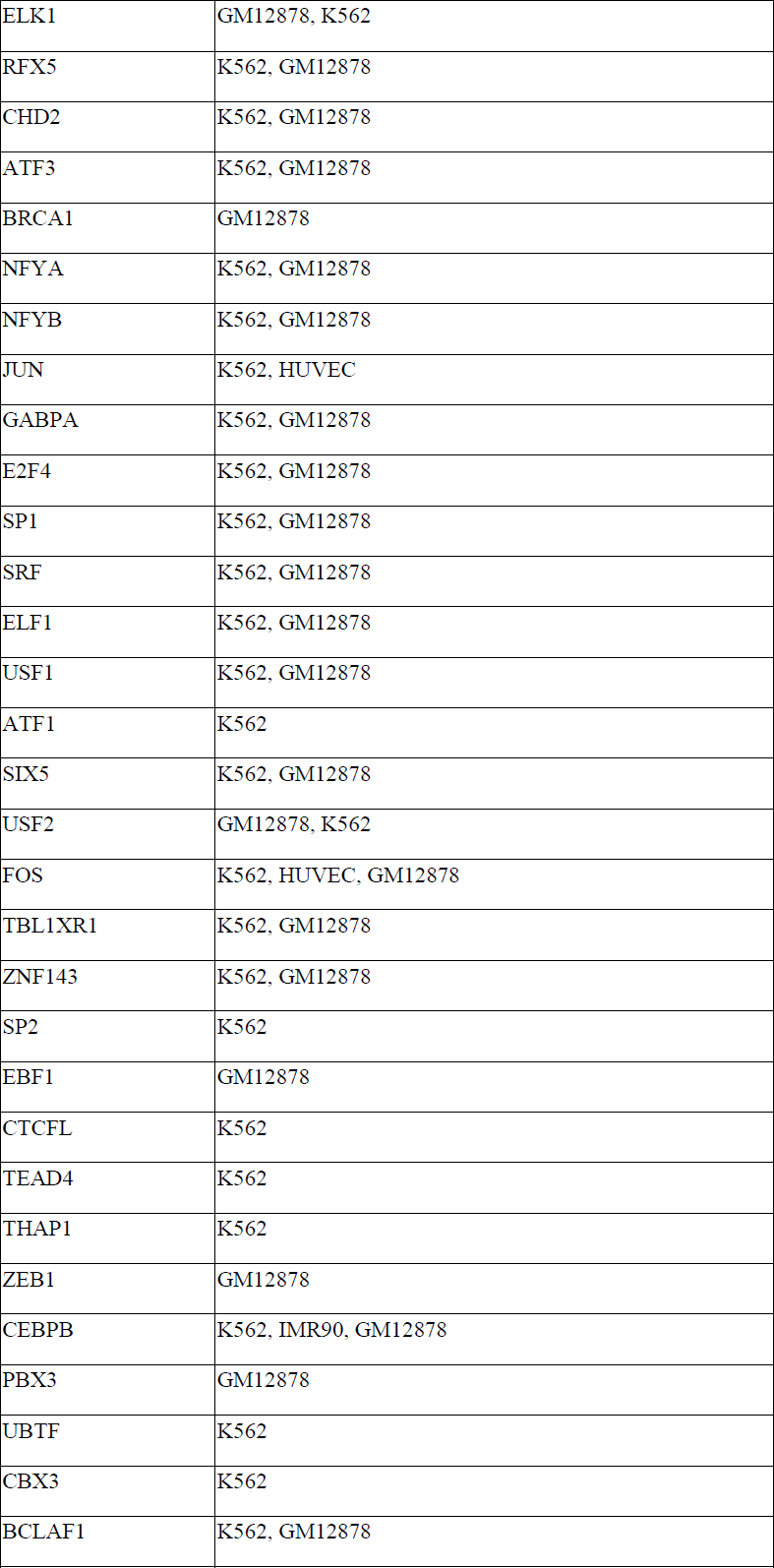
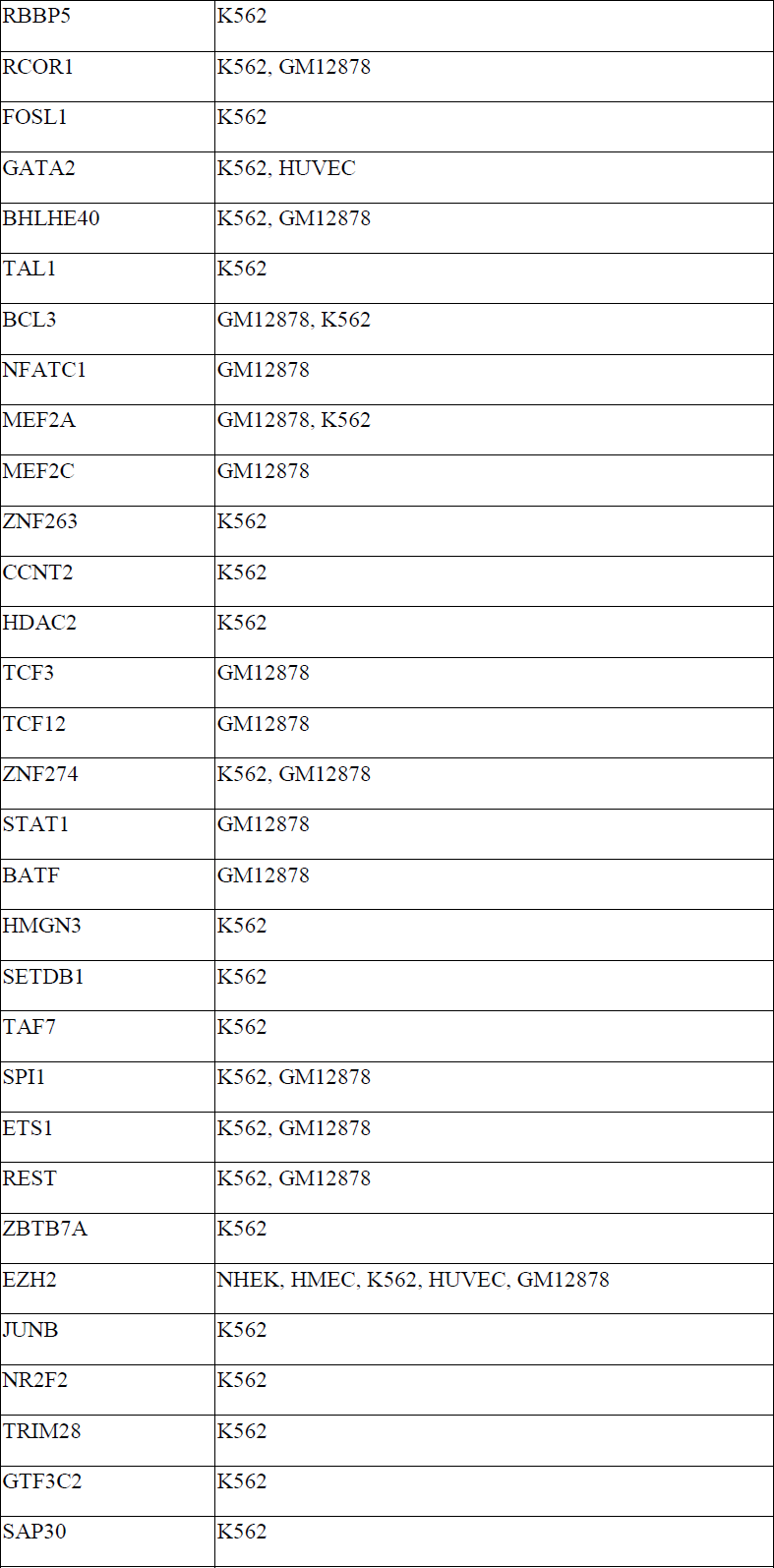
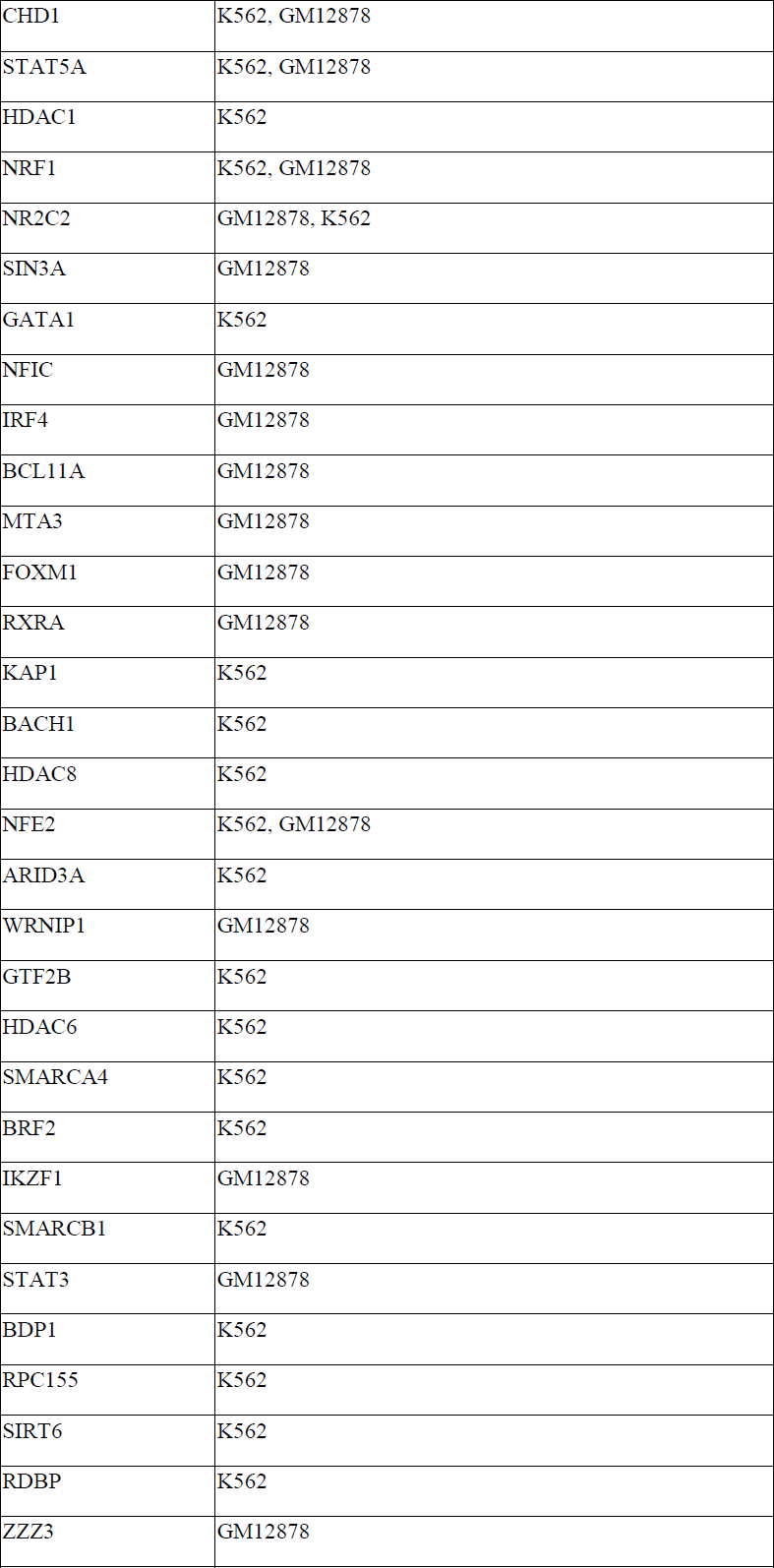
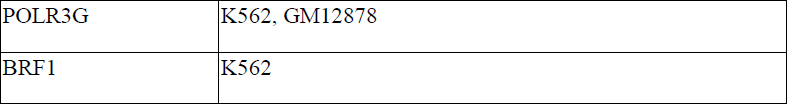
ENCODE ChIP-seq data included in our analyses (122 TFs profiled in cell lines with Hi-C data)

**Supplementary Table 3.** Enrichment of CTCF binding sites for the A compartmentalization.

| Cell line | Observed<br>A | Observed<br>B | total | Expected<br>A | Expected<br>B | D | -log 10 p-value |
| --- | --- | --- | --- | --- | --- | --- | --- |
| K562 | 48,141 | 15,524 | 63,665 | 32,056 | 31,599 | 3.06 | > 300 |
| HUVEC | 35,329 | 12,544 | 47,873 | 24,216 | 23,656 | 2.75 | > 300 |
| NHEK | 45,665 | 16,701 | 62,366 | 32,700 | 29,665 | 2.48 | > 300 |
| HMEC | 36,469 | 18,135 | 54,604 | 26,322 | 28,281 | 2.16 | > 300 |
| IMR90 | 29,184 | 14,880 | 44,064 | 21,121 | 22,942 | 2.13 | > 300 |
| GM12878 | 42,295 | 17,575 | 59,870 | 28,970 | 30,899 | 2.57 | > 300 |

**Supplementary Table 4A.** Common and cell-type specific H3K9ac sites in the comparison between GM12878 and NHEK cell lines

|  | AA | AB | BA | BB | total | R | -log10 p-value |
| --- | --- | --- | --- | --- | --- | --- | --- |
| **HMEC vs. GM12878** |  |  |  |  |  |  |  |
| **BSs only in Cell1** | 17789 | **6309** | 1708 | 5087 | 30893 | 4.48 | > 300 |
| **BSs only in Cell2** | 15292 | 567 | **3880** | 1779 | 21518 |  |  |
| **Common BSs** | 21558 | 1201 | 2319 | 3613 | 28691 |  |  |
| **HUVEC vs. GM12878** |  |  |  |  |  |  |  |
| **BSs only in Cell1** | 16635 | **6747** | 651 | 3805 | 27838 | 7.64 | > 300 |
| **BSs only in Cell2** | 16046 | 612 | **2900** | 1844 | 21402 |  |  |
| **Common BSs** | 20606 | 1173 | 966 | 2614 | 25359 |  |  |
| **HUVEC vs. HMEC** |  |  |  |  |  |  |  |
| **BSs only in Cell1** | 14992 | **4292** | 468 | 3168 | 22920 | 4.16 | > 300 |
| **BSs only in Cell2** | 16370 | 1369 | **3341** | 4301 | 25381 |  |  |
| **Common BSs** | 27220 | 2629 | 984 | 4031 | 34864 |  |  |
| **K562 vs. GM12878** |  |  |  |  |  |  |  |
| **BSs only in Cell1** | 21793 | **4294** | 1019 | 3916 | 31022 | 4.56 | > 300 |
| **BSs only in Cell2** | 14158 | 580 | **2994** | 1849 | 19581 |  |  |
| **Common BSs** | 24463 | 1239 | 1350 | 2762 | 29814 |  |  |
| **K562 vs. HMEC** |  |  |  |  |  |  |  |
| **BSs only in Cell1** | 21915 | **5083** | 982 | 3672 | 31652 | 4.15 | > 300 |
| **BSs only in Cell2** | 17015 | 1667 | **5907** | 5110 | 29699 |  |  |
| **Common BSs** | 23268 | 2498 | 1285 | 3435 | 30486 |  |  |
| **K562 vs. HUVEC** |  |  |  |  |  |  |  |
| **BSs only in Cell1** | 22645 | **3551** | 1140 | 3770 | 31106 | 5.37 | > 300 |
| **BSs only in Cell2** | 15827 | 627 | **5944** | 3772 | 26170 |  |  |
| **Common BSs** | 22627 | 1193 | 1300 | 2594 | 27714 |  |  |
| **NHEK vs. GM12878** |  |  |  |  |  |  |  |
| **BSs only in Cell1** | 19997 | **5949** | 1401 | 4154 | 31501 | 4.54 | > 300 |
| **BSs only in Cell2** | 14695 | 596 | **3111** | 1708 | 20110 |  |  |
| **Common BSs** | 21362 | 1192 | 2042 | 3023 | 27619 |  |  |
| **NHEK vs. HMEC** |  |  |  |  |  |  |  |
| **BSs only in Cell1** | 11680 | **1082** | 471 | 2402 | 15635 | 1.52 | 26 |
| **BSs only in Cell2** | 7559 | 688 | 678 | 2879 | 11804 |  |  |
| **Common BSs** | 42741 | 2511 | 2267 | 7437 | 54956 |  |  |
| **NHEK vs. HUVEC** |  |  |  |  |  |  |  |
| **BSs only in Cell1** | 18223 | **3364** | 1152 | 3363 | 26102 | 3.91 | > 300 |
| **BSs only in Cell2** | 14273 | 584 | **3426** | 3056 | 21339 |  |  |
| **Common BSs** | 26132 | 1241 | 2646 | 3560 | 33579 |  |  |
| **NHEK vs. K562** |  |  |  |  |  |  |  |
| **BSs only in Cell1** | 18782 | **5447** | 1599 | 3903 | 29731 | 3.46 | > 300 |
| **BSs only in Cell2** | 21127 | 1132 | **3998** | 3561 | 29818 |  |  |
| **Common BSs** | 23818 | 1357 | 2117 | 3026 | 30318 |  |  |

**Supplementary Table 4B.** Common and cell-type specific H3K27me3 sites in the comparison between GM12878 and NHEK cell lines

|  | AA | AB | BA | BB | total | R | p-value |
| --- | --- | --- | --- | --- | --- | --- | --- |
| **GM12878 vs. NHEK** |  |  |  |  |  |  |  |
| **BSs only in Cell1** | 7244 | 1099 | **2136** | 3306 | 13785 | 0.73 | 1.16E-18 |
| **BSs only in Cell2** | 4369 | 1026 | 1215 | 3272 | 9882 |  |  |
| **Common BSs** | 350 | 71 | 62 | 288 | 771 |  |  |
| **GM12878 vs. K562** |  |  |  |  |  |  |  |
| **MCF7 NHEK** |  |  |  |  |  |  |  |
| **BSs only in Cell1** | 5682 | 1460 | **4114** | 9381 | 20637 | 0.45 | 8.56E-113 |
| **BSs only in Cell2** | 4902 | **1066** | 904 | 3355 | 10227 |  |  |
| **Common BSs** | 129 | 64 | 55 | 168 | 416 |  |  |
| **MCF7 vs. GM12878** |  |  |  |  |  |  |  |
| **BSs only in Cell1** | 5213 | 1751 | **3637** | 9712 | 20313 | 0.54 | 2.04E-122 |
| **BSs only in Cell2** | 7176 | **1765** | 1145 | 3608 | 13694 |  |  |
| **Common BSs** | 318 | 95 | 118 | 317 | 848 |  |  |

**Supplementary Table 5.**
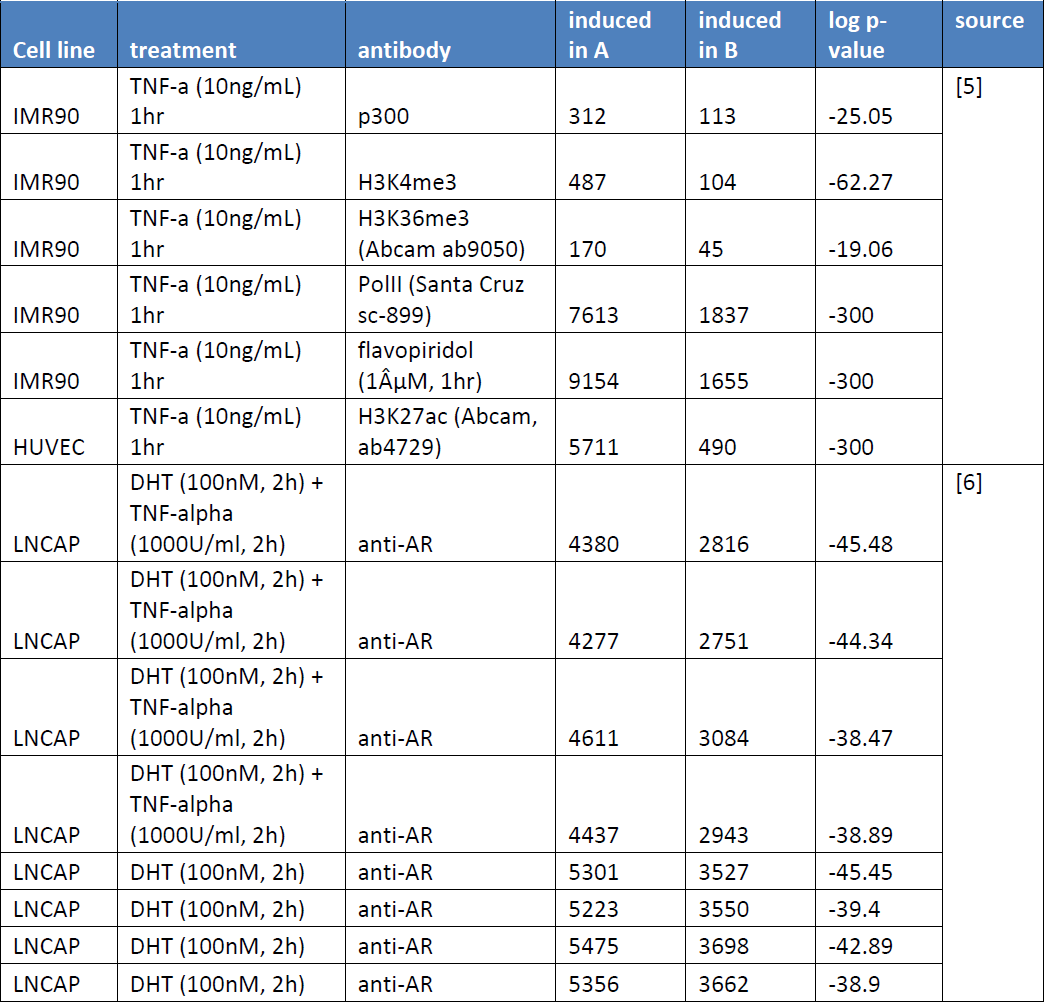
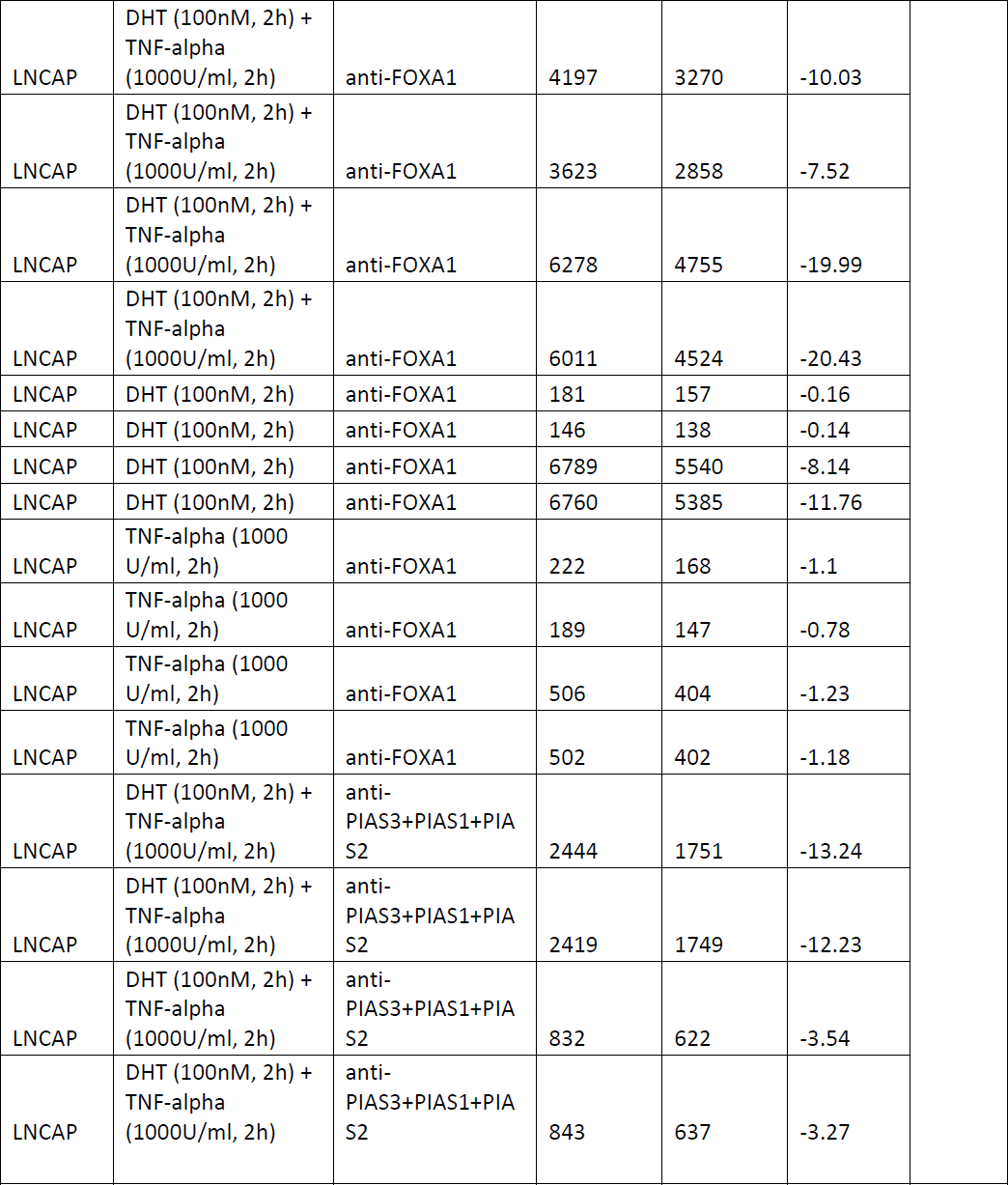
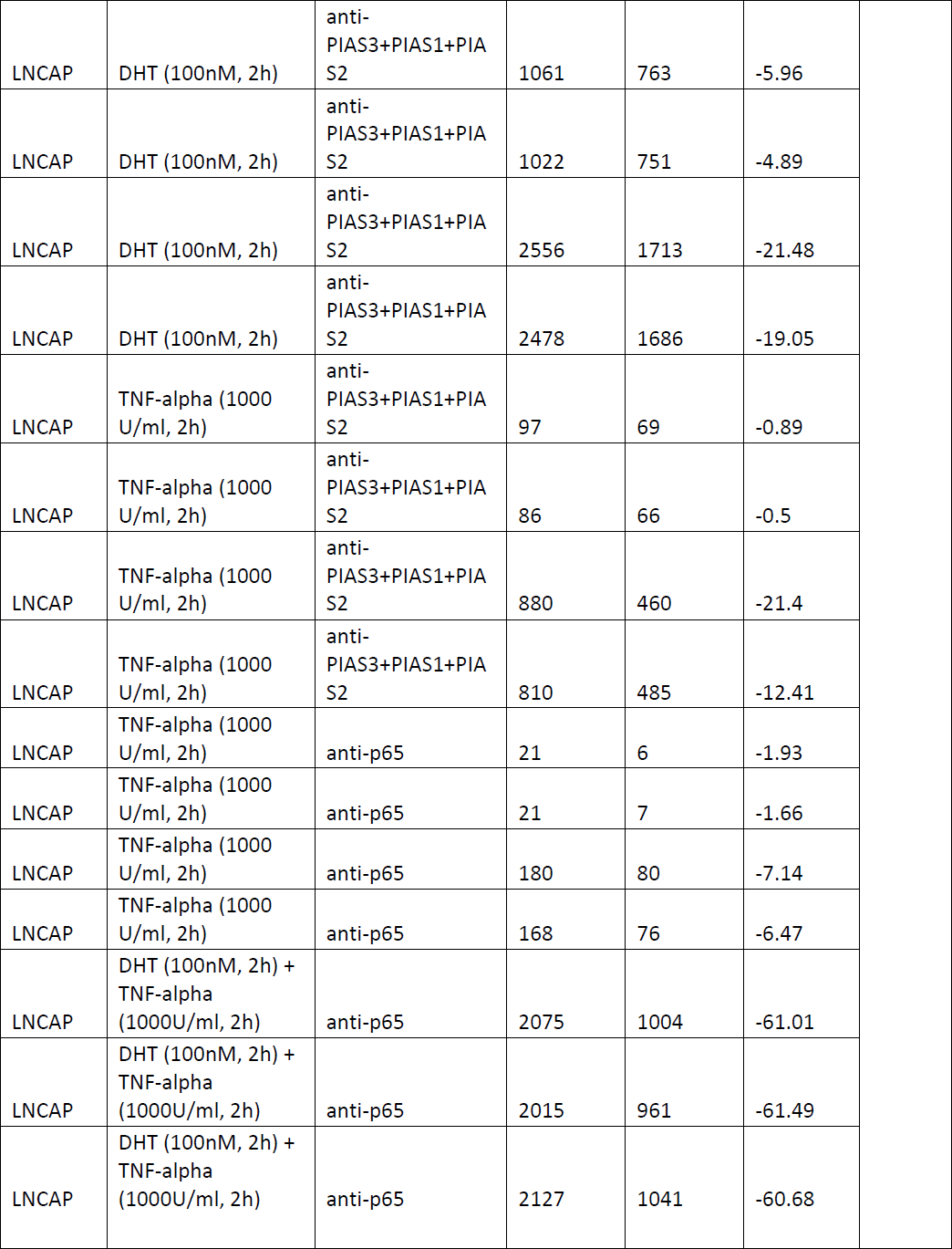
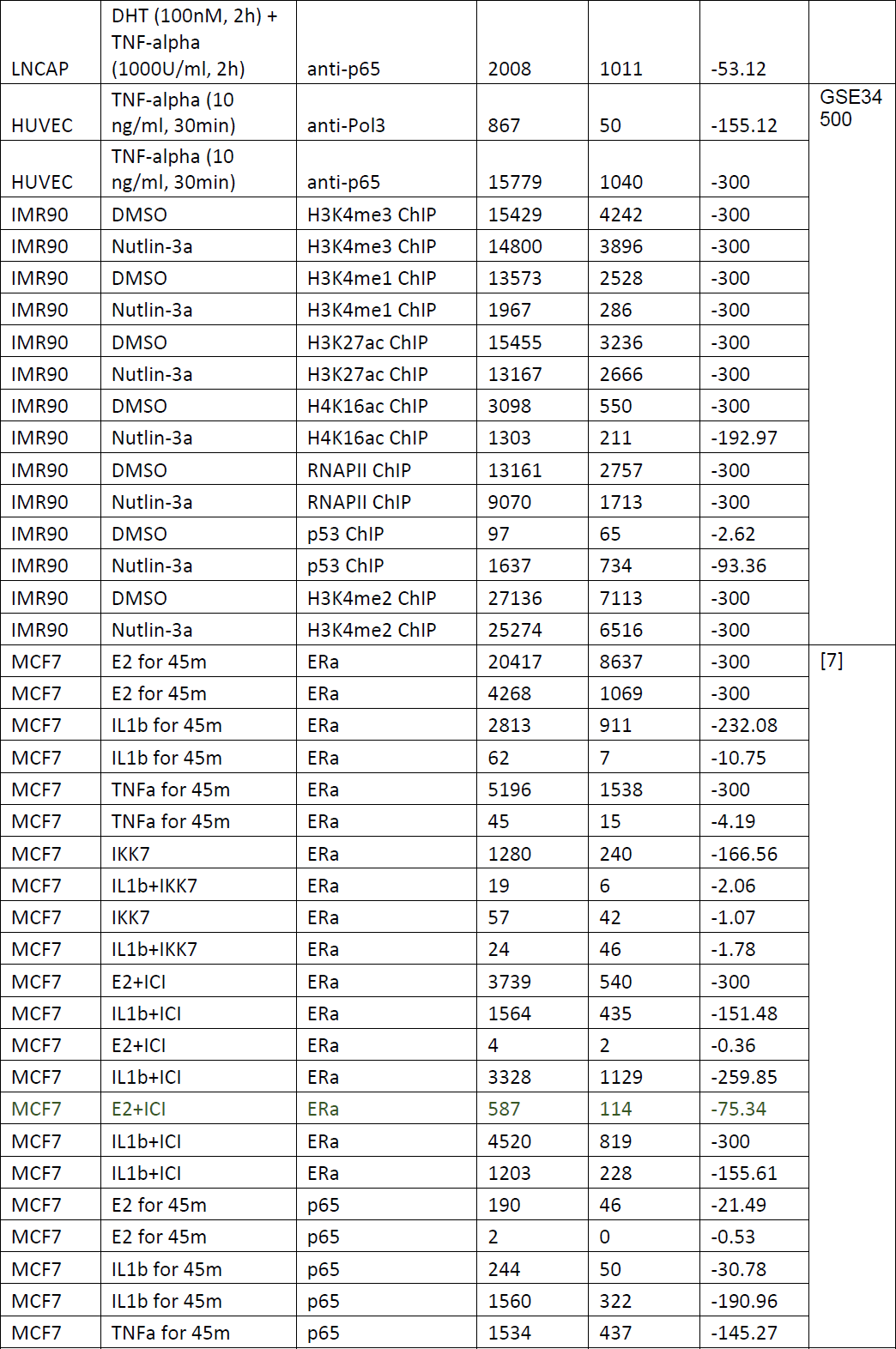
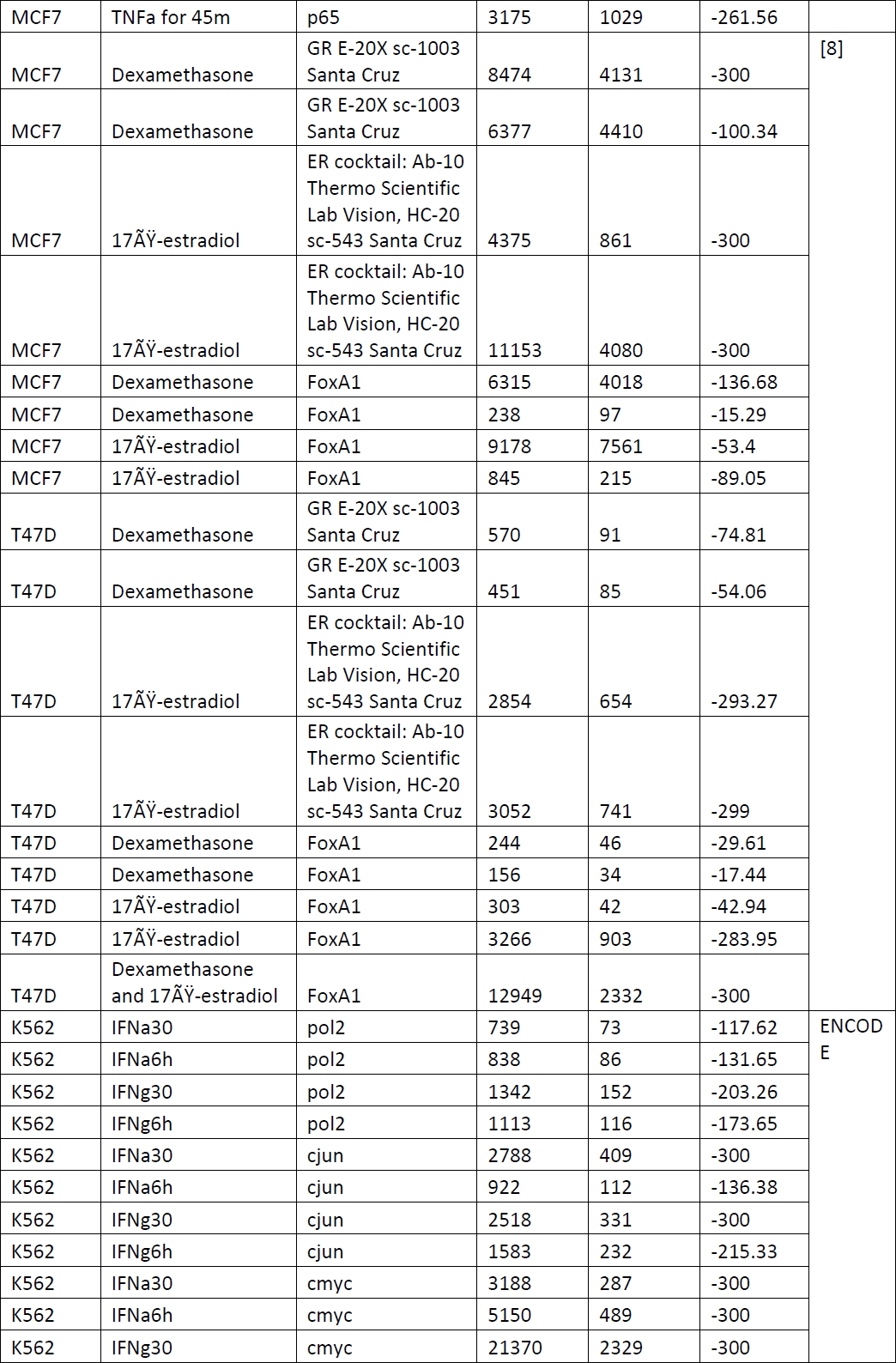
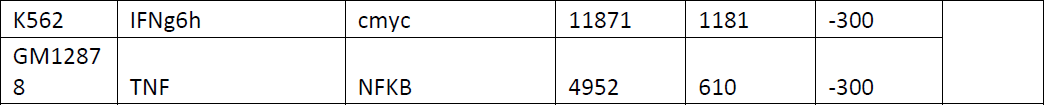
Preference of induced TF binding sites and epigenetic marks to the A compartment

**Supplementary Table 6.** Binding site induction and compartmentalization in two cell lines under the same treatment, for a particular TF.

|  | AA | AB | BA | BB | total | A/B<br>Enrichment | R | p-value |
| --- | --- | --- | --- | --- | --- | --- | --- | --- |
| **MCF7 - T47D;** |  |  |  |  |  |  |  |  |
| **Treatment: Estradiol** |  |  |  |  |  |  |  |  |
| **Antibody: ER [8]** |  |  |  |  |  |  |  |  |
| *Replicate 1* |  |  |  |  |  |  |  |  |
| Cell1_only_BSs | 2354 | 522 | 230 | 534 | 3640 | 2.33 | 1.99 | 7.06E-23 |
| Cell2_only_BSs | 1177 | 149 | 233 | 295 | 1854 | 1.63 |  |  |
| Common BSs | 1344 | 158 | 103 | 144 | 1749 |  |  |  |
| *Replicate 2* |  |  |  |  |  |  |  |  |
| Cell1_only_BSs | 7302 | 1834 | 1229 | 2751 | 13116 | 1.56 | 1.5 | 6.41E-13 |
| Cell2_only_BSs | 854 | 108 | 177 | 180 | 1319 | 1.63 |  |  |
| Common BSs | 1794 | 248 | 227 | 317 | 2586 |  |  |  |
| *Two rep combined* |  |  |  |  |  |  |  |  |
| Cell1_only_BSs | 9656 | 2356 | 1459 | 3285 | 16756 | 1.61 | 1.61 | 3.29E-29 |
| Cell2_only_BSs | 2031 | 257 | 410 | 475 | 3173 | 1.6 |  |  |
| Common BSs | 3138 | 406 | 330 | 461 | 4335 |  |  |  |
| **LNCAP - MCF7;** |  |  |  |  |  |  |  |  |
| **Treatment: TNFa;** |  |  |  |  |  |  |  |  |
| **Antibody: p65 [6]** |  |  |  |  |  |  |  |  |
| Cell1_only_BSs | 74 | 28 | 16 | 43 | 161 | 1.7 | 1.59 | 0.0014 |
| Cell2_only_BSs | 1194 | 166 | 262 | 271 | 1893 | 1.56 |  |  |
| Common BSs | 67 | 11 | 12 | 12 | 102 |  |  |  |
| **HUVEC - MCF7;** |  |  |  |  |  |  |  |  |
| **Treatment: TNFa;** |  |  |  |  |  |  |  |  |
| **Antibody: p65. Genome-wide maps of RNA Polymerase II and p65 localization in HUVECs stimulated with TNF alpha (GSE34500)** |  |  |  |  |  |  |  |  |
| Cell1_only_BSs | 12443 | 2568 | 321 | 842 | 16174 | 8 | 6.47 | 4.70E-99 |
| Cell2_only_BSs | 671 | 100 | 154 | 227 | 1152 | 1.44 |  |  |
| Common BSs | 690 | 83 | 34 | 54 | 861 |  |  |  |
| **LNCAP – HUVEC;** |  |  |  |  |  |  |  |  |
| **Treatment: TNFa;** |  |  |  |  |  |  |  |  |
| **Antibody: p65 [6]** |  |  |  |  |  |  |  |  |
| Cell1_only_BSs | 75 | 39 | 28 | 39 | 181 | 1.47 | 8.73 | 1.62E-37 |
| Cell2_only_BSs | 12237 | 374 | 3470 | 866 | 16947 | 10 |  |  |
| Common BSs | 60 | 7 | 12 | 4 | 83 |  |  |  |

**Supplementary Table 7.**
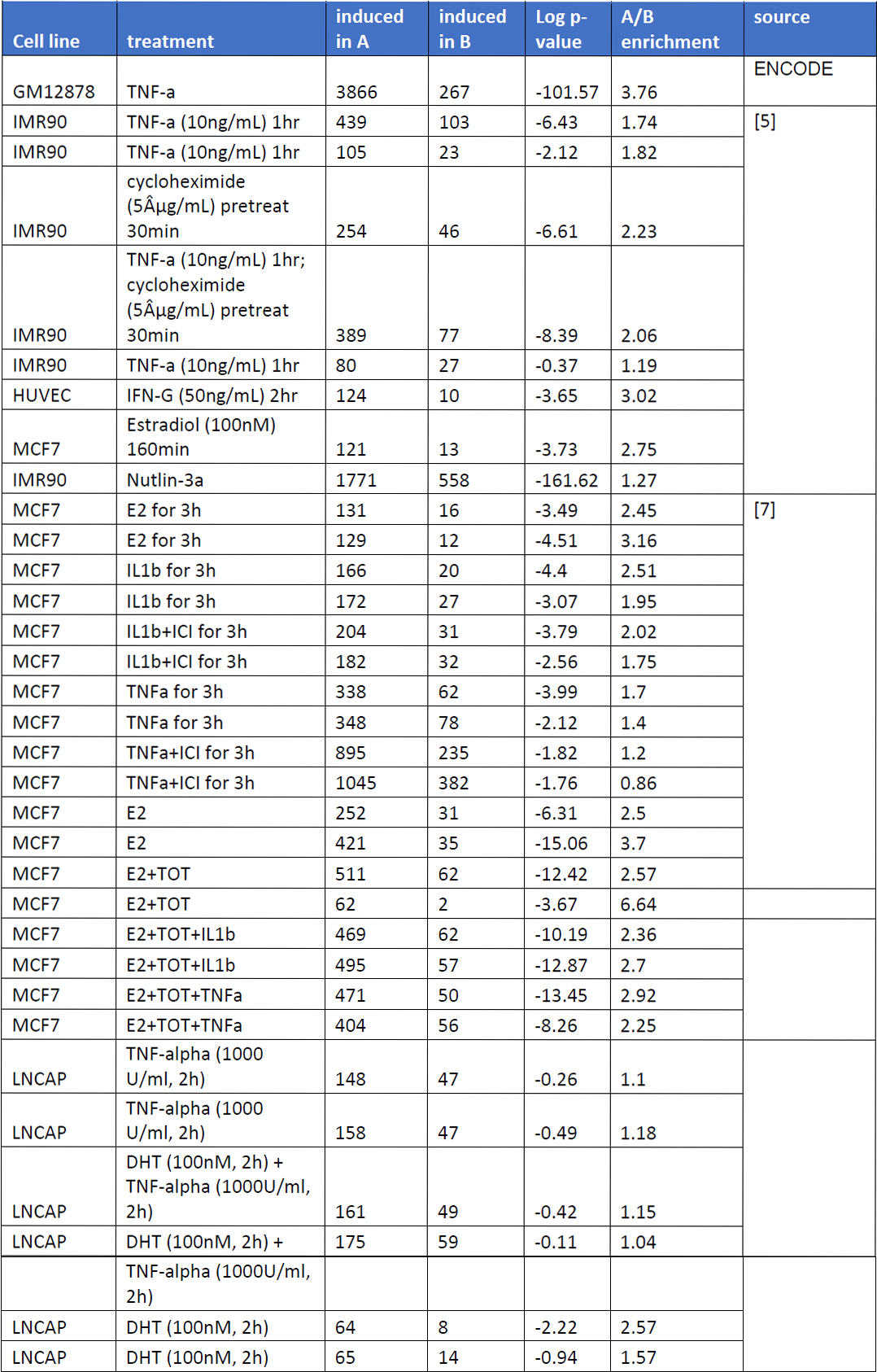
Preference of induced genes to the A compartment

**Supplementary Table 8.** Promoters of induced genes are involved, in basal condition, in higher numbers of chromatin interactions.

| Cell line | Treatment | Data type | Mean number of promoter interactions in the positive set | Mean number of promoter interactions in random sets | p-val |
| --- | --- | --- | --- | --- | --- |
| HUVEC | IFN | Hi-C | 1.46 | 0.92 | <10E-04 |
| HUVEC | TNFa | Hi-C | 1.57 | 0.95 | <10E-04 |
| K562 | SAHA | ChIA-PET | 5.44 | 3.34 | <10E-04 |
| K562 | SAHA | Hi-C | 1.12 | 0.91 | <10E-04 |
| K562 | NaBut | ChIA-PET | 5.27 | 3.37 | <10E-04 |
| K562 | NaBut | Hi-C | 1.03 | 0.92 | 0.0047 |
| HMEC | TNFa | Hi-C | 1.4 | 0.86 | <10E-04 |
| MCF7 | IL1B | ChIA-PET | 7.91 | 4.52 | <10E-04 |
| MCF7 | E2 + ICI | ChIA-PET | 4.75 | 4.53 | 0.41 |
| MCF7 | E2+TNFa | ChIA-PET | 7.16 | 4.53 | <10E-04 |
| MCF7 | E2 | ChIA-PET | 4.94 | 4.53 | 0.04 |
| MCF7 | E2+IL1b | ChIA-PET | 7.51 | 4.52 | <10E-04 |
| MCF7 | E2 | ChIA-PET | 8.16 | 4.53 | <10E-04 |
| MCF7 | IL1b+ICI | ChIA-PET | 7.67 | 4.52 | <10E-04 |
| MCF7 | TNFa | ChIA-PET | 7.27 | 4.52 | <10E-04 |
| MCF7 | E2 | ChIA-PET | 10.63 | 4.53 | <10E-04 |
| MCF7 | TNFa+ICI | ChIA-PET | 6.89 | 4.53 | <10E-04 |
| IMR90 | Nutlin-3a | Hi-C | 1.21 | 0.86 | <10E-04 |
| IMR90 | TNFa | Hi-C | 1.25 | 0.86 | <10E-04 |
| IMR90 | TNFa+cycloheximide | Hi-C | 1.24 | 0.86 | <10E-04 |
| GM12878 | TNFa | ChIA-PET | 3.91 | 1.81 | <10E-04 |
| GM12878 | TNFa | Hi-C | 1.77 | 1.27 | <10E-04 |

**Supplementary Table 9.**
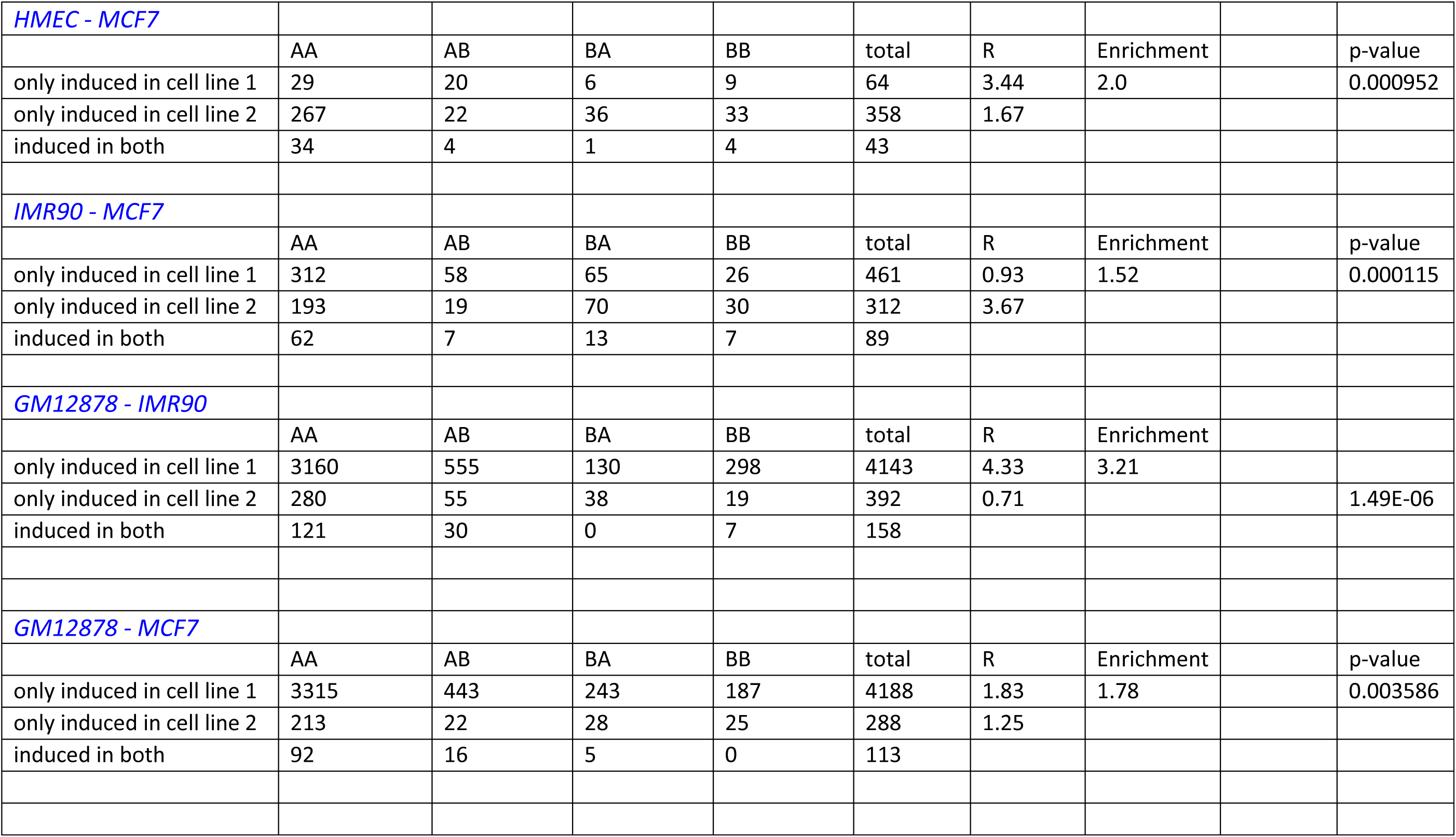
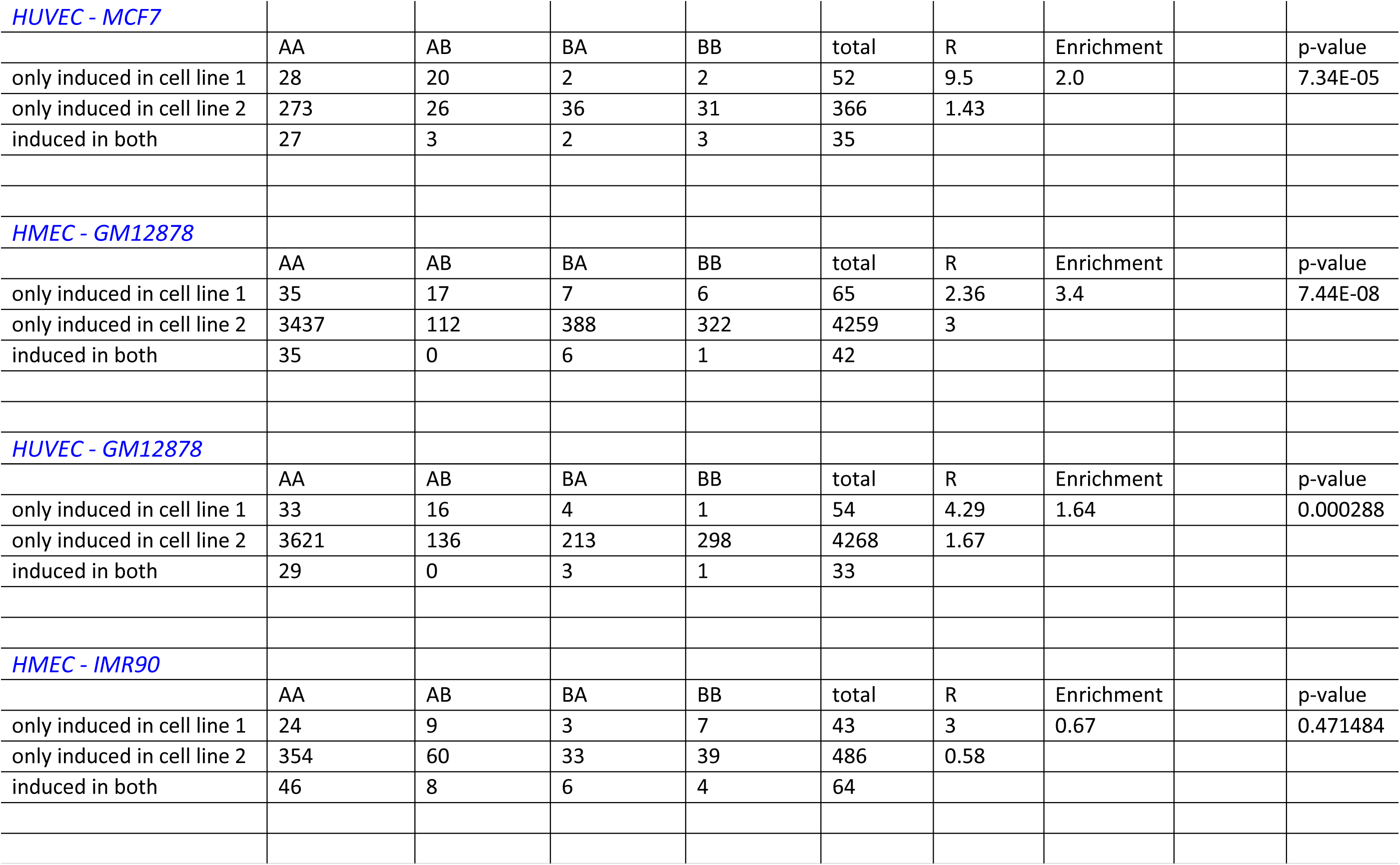
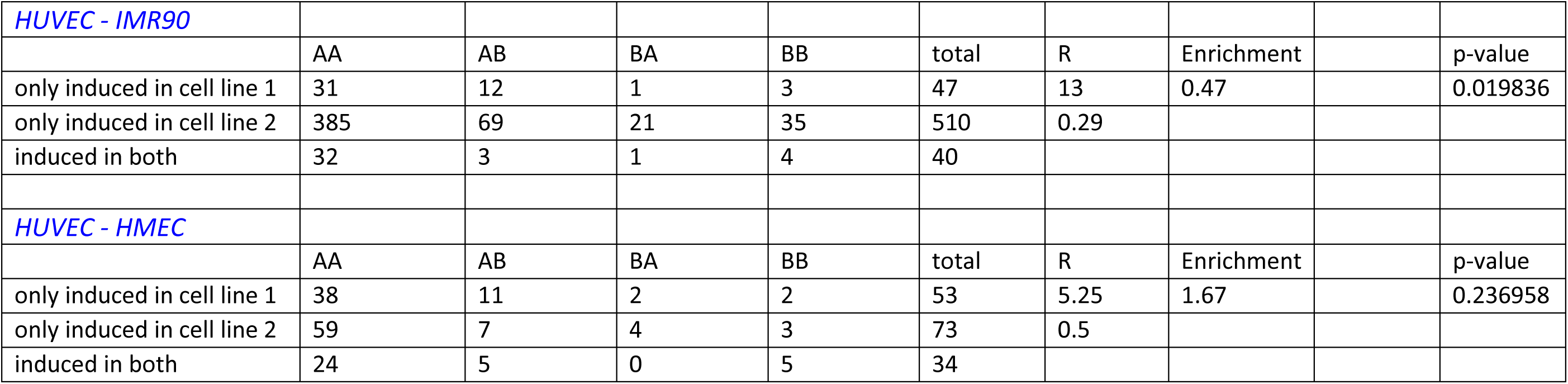
Preference of cell-type specific induced genes to cell-type specific A compartment

**Supplementary Figure 1.**
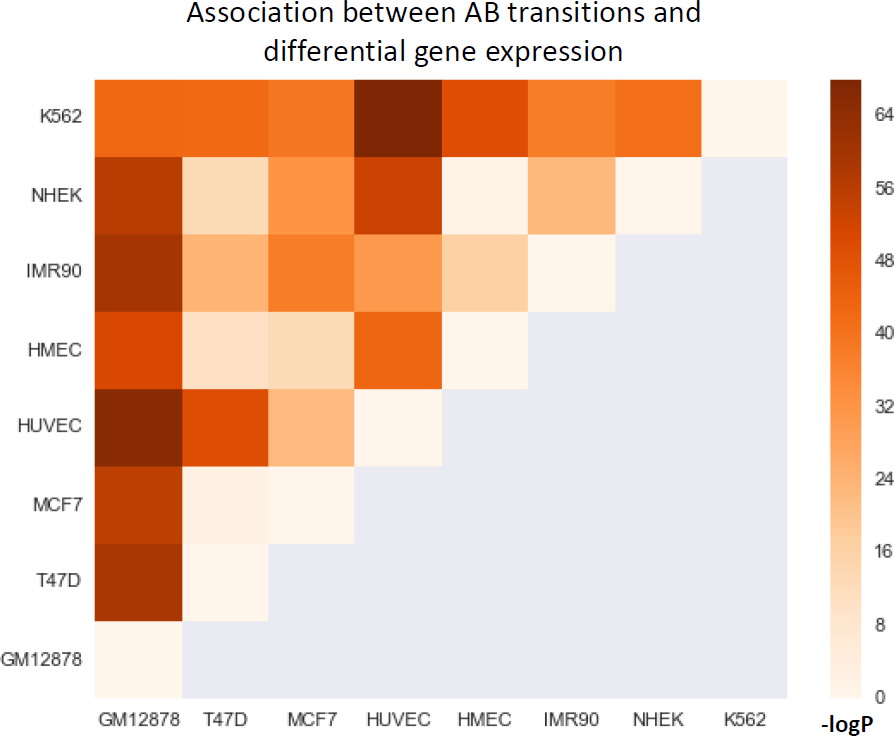
Association between changes in A/B compartmentalization and differential gene expression between cell types. For each pair of cell lines, we examined the difference (fold-change) in expression level between genes assigned to the AB and BA sets (for a pair of cell lines 1 and 2, AB: genes located in the A compartments in cell line 1 and in B in cell line 2; BA: genes located in the B compartment in cell line 1 and in A in cell line 2). For 27 out of 28 pairwise comparisons (all except HMEC-NHEK), we observed a highly significant association (FDR<<5%) between differential compartmentalization and expression. (p-values calculated using Wilcoxon’s test.)

**Supplementary Figure 2.**
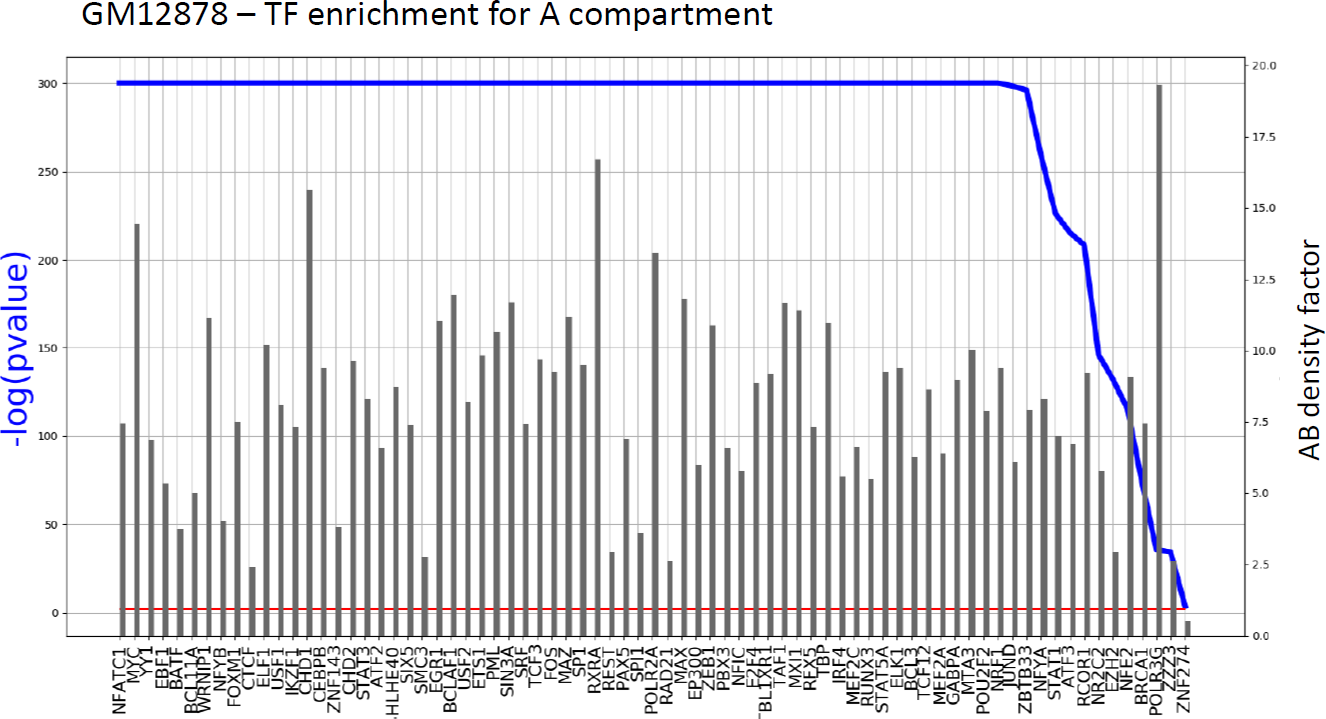
Enrichment of TFBSs in the A compartment. ChIP-seq experiments are sorted by p-value, A-B density factors are represented by bars. Red line indicates p-value = 0.01. Shown are experiments in the GM12878 cell line. Similar results were observed for all other cell lines (data not shown).

**Supplementary Figure 3.**
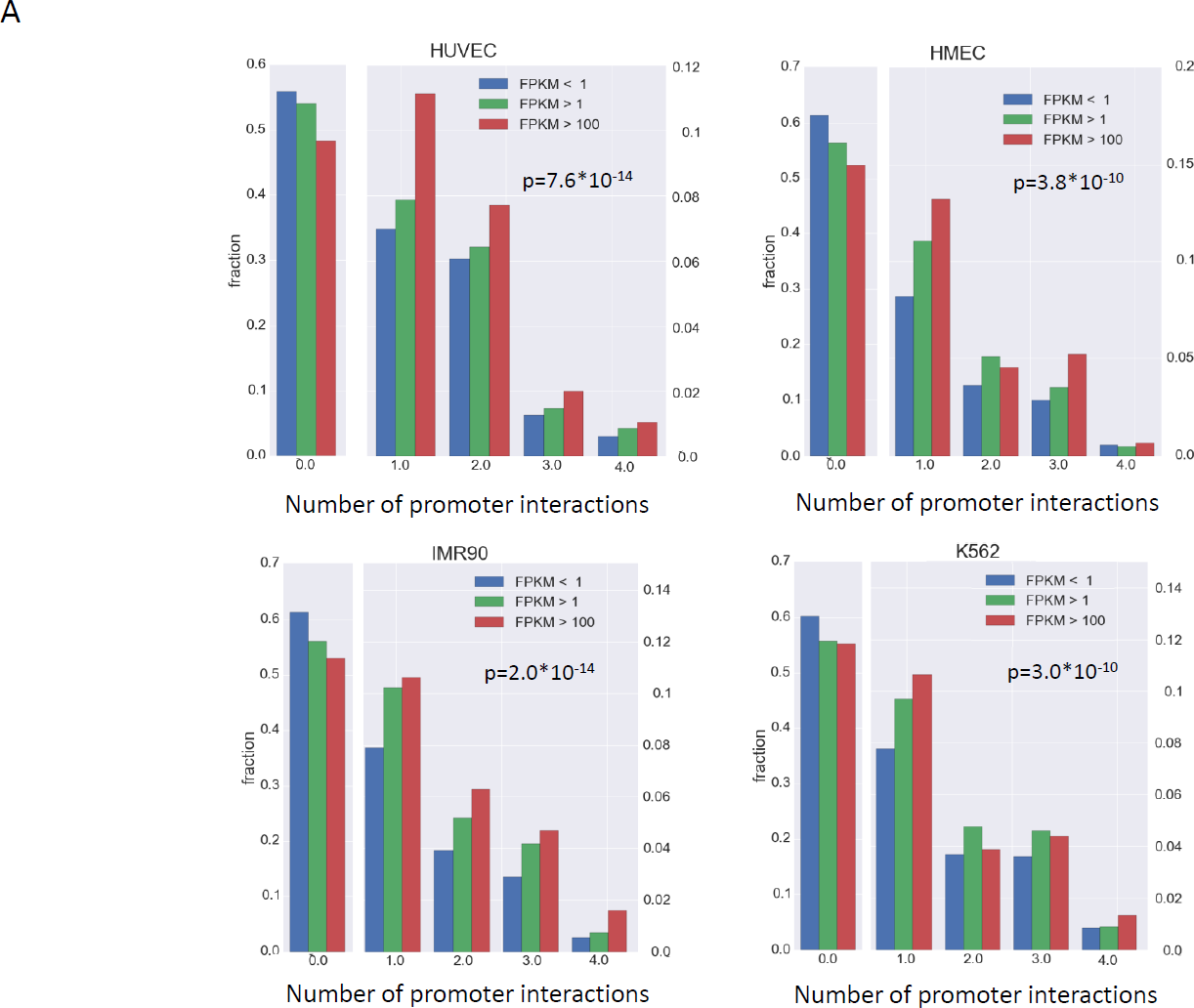
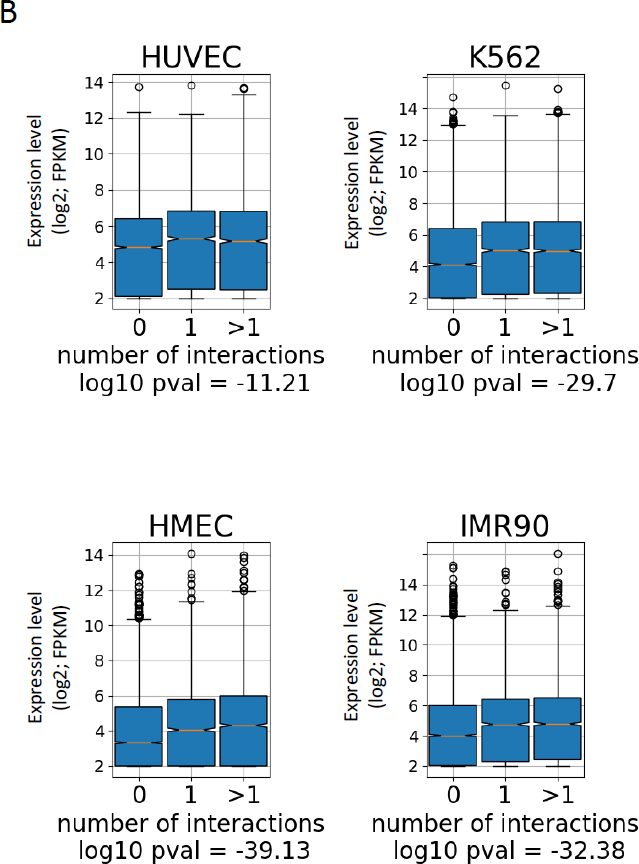
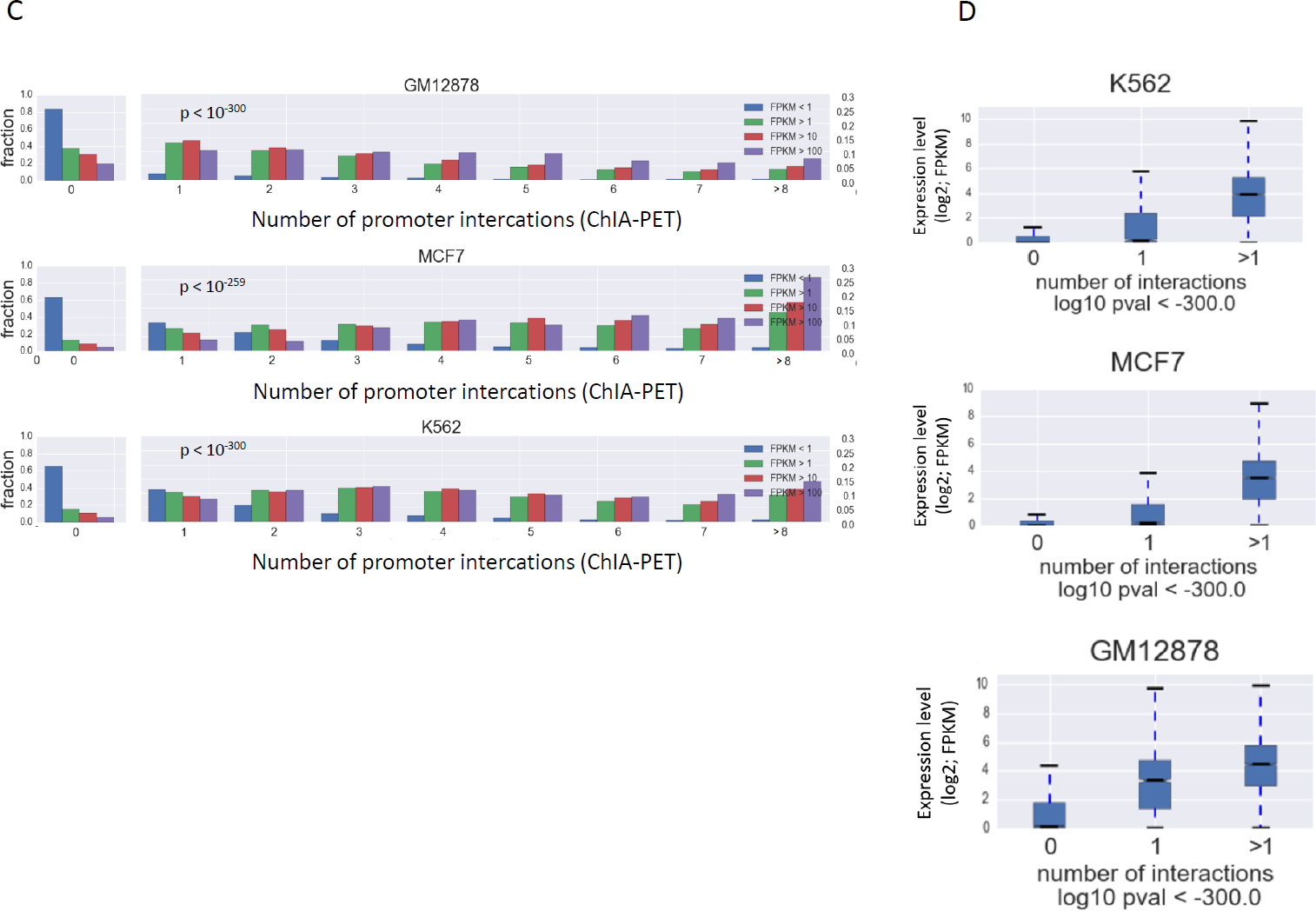
Gene expression levels vs. promoter interactions in compartment. **A**. The same analyses described in the legend of Fig. 3 are applied here to additional cell lines.

